# Chloroplasts alter their morphology and accumulate at the pathogen interface during infection by *Phytophthora infestans*

**DOI:** 10.1101/516443

**Authors:** Zachary Savage, Cian Duggan, Alexia Toufexi, Pooja Pandey, Yuxi Liang, María Eugenia Segretin, Lok Him Yuen, David C. A. Gaboriau, Alexandre Y. Leary, Yasin Tumtas, Virendrasinh Khandare, Andrew D. Ward, Stanley W. Botchway, Benji C. Bateman, Indranil Pan, Martin Schattat, Imogen Sparkes, Tolga O. Bozkurt

## Abstract

Upon immune activation, chloroplasts switch off photosynthesis, produce anti-microbial compounds, and associate with the nucleus through tubular extensions called stromules. Although it is well-established that chloroplasts alter their position in response to light, little is known about the dynamics of chloroplasts movement in response to pathogen attack. Here, we report that chloroplasts accumulate at the pathogen interface during infection by the Irish potato famine pathogen *Phytophthora infestans*, associating with the specialized membrane that engulfs the pathogen haustorium. Chemical inhibition of actin polymerization reduces the accumulation of chloroplasts at the pathogen haustoria, suggesting this process is partially dependent on the actin cytoskeleton. However, chloroplast accumulation at haustoria does not necessarily rely on movement of the nucleus to this interface and is not affected by light conditions. Stromules are typically induced during infection, embracing haustoria and interconnecting chloroplasts, to form dynamic organelle clusters. We found that infection-triggered stromule formation relies on BRASSINOSTEROID INSENSITIVE 1-ASSOCIATED KINASE 1 (BAK1) mediated surface immune signaling, whereas chloroplast repositioning towards haustoria does not. Consistent with the defense-related induction of stromules, effector mediated suppression of BAK1 mediated immune signaling reduced stromule formation during infection. On the other hand, immune recognition of the same effector stimulated stromules, presumably via a different pathway. These findings implicate chloroplasts in a polarized response upon pathogen attack and point to more complex functions of these organelles in plant-pathogen interactions.

## Introduction

*Phytophthora infestans* is an oomycete pathogen that causes potato late blight, one of the most historically important and economically devastating crop diseases. The pathogen penetrates host cells via haustoria, infection structures that extend from its intercellular invasive hyphae. Haustoria are surrounded by the plant-derived extrahaustorial membrane (EHM), across which effectors secreted by the pathogen translocate inside the host cell (Wang et al., 2017; Whisson et al., 2007, 2016). This interface is key to the success or failure of infection and is therefore targeted by focal immune responses of the plant (Bozkurt et al., 2011; Dagdas et al., 2018; Kwon et al., 2008). This includes the deposition of callose, redirection of autophagy, and movement of the nucleus towards the site of penetration (Dagdas et al., 2018; Griffis et al., 2014; Jones & Dangl, 2006). While continuous with the plasma membrane, there is a stark difference in the biochemical composition of the EHM and the plasma membrane (Bozkurt et al., 2014, 2015).

The EHM typically lacks the surface localized pattern recognition receptors (PRRs), which activate downstream immune responses through recognition of pathogen associated molecular patterns (PAMPs) (Bozkurt et al., 2014, 2015). Once PPRs detect PAMPs, downstream signaling is triggered, often in co-ordination with a co-receptor such as BRASSINOSTEROID INSENSITIVE 1-ASSOCIATED RECEPTOR KINASE1 (BAK1), to induce an immune response (Chaparro-Garcia et al., 2011; Heese et al., 2007). To counteract this, pathogens typically deploy host-translocated effectors to subvert surface mediated immunity. For example, *P. infestans* host-translocated RXLR effector AVR3a suppresses BAK1 mediated surface immune responses (Chaparro-Garcia et al., 2015). However, presumably, plant basal responses still contribute to immunity against adapted pathogens, as immune suppression by effectors is often partial, as in the case of AVR3a (Chaparro-Garcia et al., 2015). Moreover, downregulation of *BAK1* gene expression in solanaceous model plant *Nicotiana benthamiana* leads to significantly enhanced pathogen growth, further highlighting the importance of surface mediated immunity against *P. infestans* (Chaparro-Garcia et al., 2015). This makes *N. benthamiana* an excellent model for studies to dissect the functional principles of basal plant immunity against *P. infestans*. Furthermore*, N. benthamiana* lacks the specialized nucleotide-binding leucine-rich repeat (NLR) type of immune receptors that can sense *P. infestans* effectors intracellularly. This is also advantageous because it allows for live cell imaging of *P. infestans* infection, as NLR mediated immunity often triggers a form of programmed cell death at the site of infection known as the hypersensitive response (Wu et al., 2017).

Activation of immunity at the cell surface stimulates chloroplasts to shut down photosynthesis, synthesize defense hormone precursors, and generate reactive oxygen species (ROS) (Padmanabhan & Dinesh-Kumar, 2010; Su et al., 2018), indicating that chloroplasts are major components of the plant defense system. Pathogens are known to target chloroplasts with effector proteins, further highlighting their importance in immunity (Jelenska et al., 2007; Pecrix et al., 2019; Petre et al., 2016; Zabala et al., 2015). Interestingly, several genes associated with resistance to oomycete pathogens were found to encode chloroplast-localized proteins (Belhaj et al., 2009; Van Damme et al., 2009). Chloroplasts also produce stroma filled tubules (stromules) in response to a range of elicitors, including phytohormones, ROS and the bacterial PAMP, flg22 (Brunkard et al., 2015; Caplan et al., 2015; Gray et al., 2012). While the exact function(s) of stromules is still unclear, they have been implicated in immunity, chloroplast movement, and connection to the plant cell nucleus (Caplan et al., 2015; Kumar et al., 2018). Immune stimulation by PAMPs and ROS also induce the association between chloroplasts and the nucleus, hinting at potential defense-related roles of chloroplast re-distribution during infection (Ding et al., 2019). However, the molecular and physiological mechanism of how chloroplast immunity is launched against invading pathogens is unclear.

Here, we used quantitative confocal microscopy to investigate the spatial dynamics of chloroplasts in living plant cells infected by *P. infestans*. We show that chloroplasts accumulate around haustoria in a dynamic fashion, but this process does not necessarily rely on movement of the nucleus towards the haustorium. We found that the actin-cytoskeleton, but not light conditions, are critical for chloroplast positioning around the haustorium. Our microscopy analyses using optical tweezers suggest association between chloroplasts and the EHM. Finally, we demonstrate that chloroplasts also alter their morphology by induction of stromules as a defense response, whereas effectors can counteract this process.

## Results

### Chloroplasts accumulate at the host-pathogen interface in an actin dependent manner but irrespective of light conditions

While the immune-related roles of chloroplasts in producing anti-microbial compounds and defense signaling molecules are well established, little is known on the subcellular dynamics of chloroplast movement and stromule induction during infection. To gain insights into this, we monitored the live infection of *N. benthamiana* by *P. infestans*, specifically targeting cells containing a pathogen haustorium (haustoriated cells). In this infection model, haustoria are generally visible in the host epidermal cells, where the chloroplasts are smaller and less abundant than in the mesophyll. We reasoned that the more sparsely distributed chloroplasts of these epidermal cells may redistribute towards the site of infection to partake in a localized intracellular immune response.

During infection of *N. benthamiana* with the fluorescently tagged strain of *P. infestans*, 88069td, haustoria are easily visible in host cells. Confocal microscopy of infected leaf epidermal cells stably expressing GFP in chloroplast stroma (CpGFP herein) revealed that chloroplasts associate with 40% of haustoria (*N* = 280 haustoria) (Fig. 1A, Video S1). Haustoria were often associated with multiple chloroplasts, and these chloroplasts were often mobile around the site of infection (Fig. 1A, Video S2).

**Figure 1:**
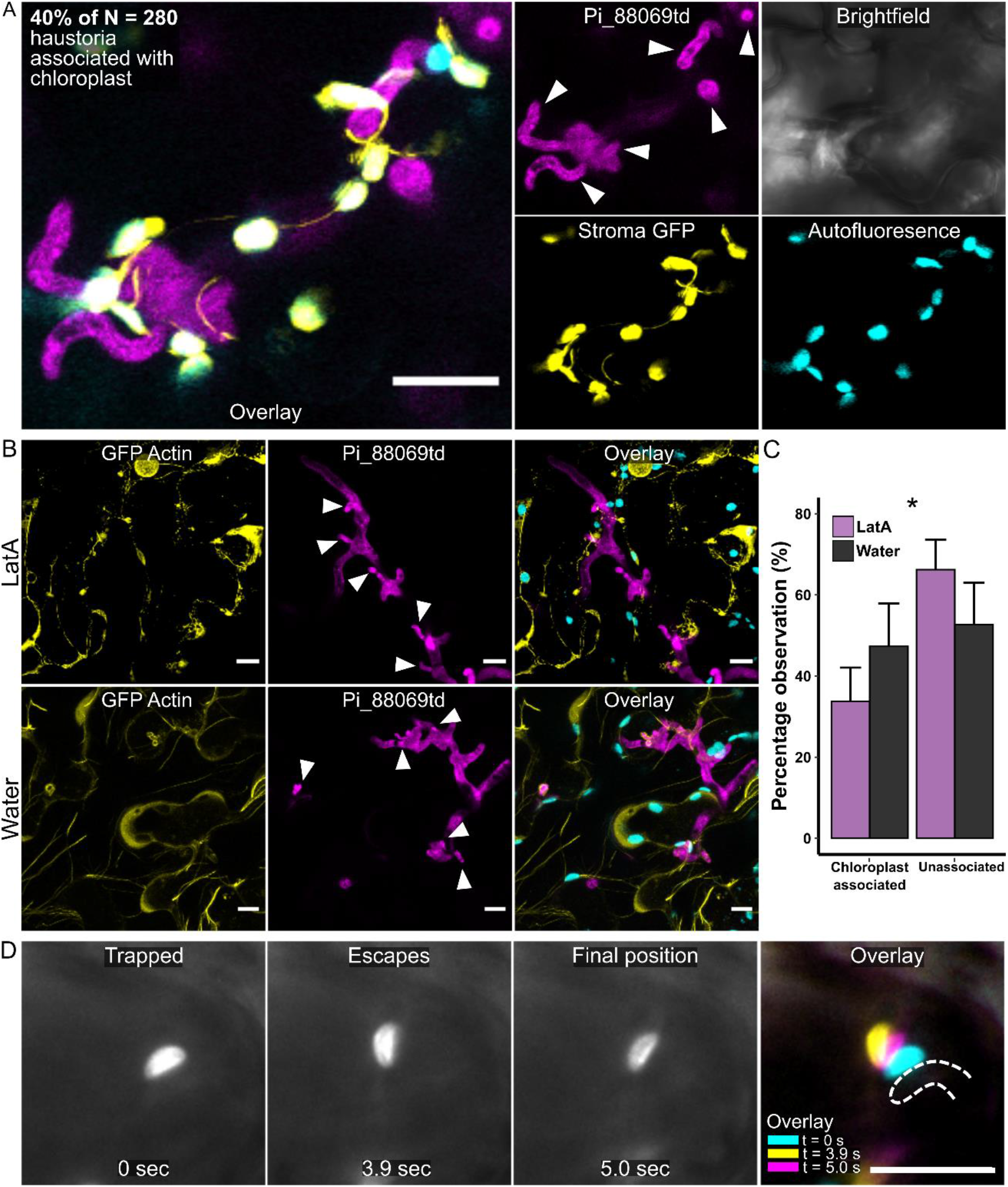
Chloroplasts accumulate at the host-pathogen interface during infection in an actin dependent manner. **(A)** Maximum projection confocal micrographs of *N. benthamiana* plants expressing GFP in chloroplast stroma (CpGFP), infected with *P. infestans* strain 88069td, showing chloroplast positioning around pathogen haustoria. Haustoria marked by white arrow heads. **(B)** Maximum projection confocal micrographs of wild-type *N. benthamiana* plants, transiently expressing GFP actin chromobody, infected with *P. infestans* strain 88069td, following 24 hr treatment with water or 1 μM of chemical actin polymerization inhibitor LatA. Haustoria marked by white arrow heads. **(C)** Bar plots showing percentage of chloroplast accumulation at haustoria following 24 hr treatment with water or 1 μM of chemical actin polymerization inhibitor LatA. Observations made in wild-type plants infected with *P. infestans* strain 88069td, across 4 separate biological replicates, totaling 148 and 93 haustoria in the LatA and water conditions respectively. Error bars show confidence intervals. Asterisk denotes *p* < 0.05 as determined by Fisher’s Exact test. **(D)** GFP channel in grayscale from TIRF microscope. Time-lapse showing laser capture of haustorium associated chloroplast in CpGFP plant where the automated trapping routine traps and attempts to move chloroplast 10 μm. Chloroplast escapes the trap at 3.9 seconds before it springs back towards the original position at 5 seconds. Dotted line shows outline of haustorium marked by RFP:REM1.3. All scale bars are 10 μm.

We investigated whether positioning of chloroplasts around haustoria was a response to infection or due to the result of chance encounter, i.e., because of haustoria coincidentally penetrating the cell at the position where there is a chloroplast. To check whether this association occurred at a greater frequency than would be expected by random chance encounter, we developed an unbiased method to position mock haustoria throughout micrographs of infected tissue (Fig. S1A). For each real haustorium in an image, a straight line was drawn from the point at which the haustorium entered the cell until the opposite edge of the same cell was reached; this end point was imagined as the position of a mock haustorial penetration. We then categorized whether each mock haustoria was immediately adjacent to, or in contact with, a chloroplast in that position (Fig. S1A). If chloroplast accumulation at haustoria was random, we would expect rates of actual and mock haustoria to be the same on opposite edges of the cell. Compared to the actual haustoria, there were significantly fewer instances contact between chloroplasts and mock haustoria (Fig. S1B), suggesting the association between haustoria and chloroplasts was not due to random encounter during cell penetration.

To gain further insights into chloroplast positioning around the haustorium, we monitored chloroplast dynamics in haustoriated cells with time lapse-microscopy. Here, we observed several instances of chloroplast movement towards the haustoria, suggesting an active relocation of chloroplasts towards the site of intracellular infection (Video S3, S4). Similar observations were made during wild-type *P. infestans* infection, visualizing the EHM with RFP:REM1.3 (Video S5). Additionally, we observed some instances where haustoria that were associated with chloroplasts collapse during the acquisition of the time-lapse (Video S6, S7). This indicates that some of the haustoria-chloroplast associations could be missed out during image quantification as it is not always possible to identify collapsed haustoria.

The cytoskeleton, and particularly actin, has known roles in chloroplast movement (Wada & Kong, 2018), nucleus movement (Higa et al., 2014), stromule interactions (Erickson et al., 2018; Kumar et al., 2018), and cell polarization towards the sites of pathogen penetration (Kobayashi & Hakuno, 2003; Opalski et al., 2005). Therefore, we next tested whether chemical inhibition of actin polymerization by Latrunculin-A (LatA) would influence the accumulation of chloroplasts at haustoria. To monitor successful disruption of actin, a GFP tagged actin chromobody was transiently expressed during 1 μM LatA treatment of infected wild-type plants by *P. infestans* 88069td (Fig. 1B). We also tested a range of LatA concentrations for ability to visually disrupt actin filaments before completing the infection microscopy, in order to use a minimal concentration of LatA (Fig. S2). Treatment with LatA significantly reduced the frequency of chloroplast-haustoria associations compared to the control condition (34% of *N* = 148 haustoria, and 47% of *N* = 93 haustoria respectively) (Fig. 1C). This points to a role for actin in the accumulation of chloroplasts at haustoria. However, we are cautious of overinterpreting this data due to the non-specific nature of the drug and its plausible ability to influence the growth and virulence of the pathogen (Ketelaar et al., 2012). By using the GFP marker of actin to ensure inhibition in the cells imaged, we sacrificed the ability to monitor stromule induction during this experiment.

Finally, we also tested whether light exposure has any impact on accumulation of chloroplasts at haustoria. For the dark condition, we kept *P. infestans* 88069td infected CpGFP leaves in the dark for 2 days prior to imaging (dark-dark), while we kept the control group in regular day-night light cycling conditions (light-dark). Live cell infection microscopy revealed no difference in chloroplast accumulation between the two lighting conditions (Fig. S3). Collectively, these findings indicate that chloroplasts actively position around haustoria during *P. infestans* infection through a process that requires actin polymerization, whereas this process is not affected by changes in light conditions.

### Optical tweezers reveal the association between chloroplasts and the haustoria

Accumulating evidence points to the importance of organelle membrane contacts in response to various physiological or stress conditions (Helle et al., 2013; Liu & Li, 2019; Silva et al., 2020). To investigate the nature of chloroplasts association with haustoria/EHM, we used optical tweezers in combination with Total Internal Fluorescence Microscopy (TIRF) in CpGFP plants, infected with the wild-type *P. infestans* 88069, and transiently expressing the extrahaustorial membrane (EHM) marker, RFP:REM1.3 (Bozkurt et al., 2014). Relocation of chloroplasts further than a 10 μm threshold was considered a successful movement by optical tweezers. This threshold was set to ensure consistency between experiments; 10 μm was chosen as this distance was small enough to avoid side effects from moving the chloroplast towards the vacuole, such as pushing the chloroplast into the tonoplast.

Using optical tweezers, we successfully trapped and moved 17% of chloroplasts (*N* = 29) in non-haustoriated cells beyond the threshold using the automated trapping routine. In comparison, we were unable to trap and move any chloroplasts (0%, *N* = 18) neighboring haustoria past 10 μm, suggesting an association or connection between the chloroplasts and the EHM. Consistent with this, we recorded instances where these chloroplasts were initially pulled away from the EHM, but before they passed 10 μm, they escaped the trap and sprang back towards their former position (22%, *N* = 18) (Fig. 1D, Video S8). These results suggest that chloroplasts may establish secure contacts with the EHM, however, further genetic and biochemical evidence supporting these findings are required to reach definitive conclusions.

### Chloroplasts can associate with haustoria independently of the host nucleus

Relocation of the plant nucleus towards pathogen penetration sites was reported as a hallmark of plant focal immune responses (Griffis et al., 2014). Given recent reports showing the association of chloroplasts with the nucleus during plant stress responses (Caplan et al., 2015; Ding et al., 2019; Erickson et al., 2018), it is possible that chloroplasts are dragged to haustorium along with the nucleus. However, this is unlikely because there is only a single nucleus per plant cell, and *P. infestans* can form multiple haustoria that are in contact with chloroplasts within a single cell (Fig. 1A).

Nevertheless, we quantified the numbers of haustoria associated with chloroplasts alone, compared to chloroplasts and nuclei together, to determine the extent to which chloroplast positioning around the haustoria correlates with the presence of nucleus. To track the nucleus, we used leaf epidermal cells stably expressing GFP in the endoplasmic reticulum (ER-GFP herein), to visualize the silhouette of the nucleus (Fig. 2). Confocal microscopy followed by manual quantification of 463 haustoria revealed that 10% were associated with both a chloroplast and the plant nucleus (Fig. 2A), while 26% were associated with just a chloroplast but not a nucleus (Fig. 2B). Very rarely (2%), a haustorium was in contact with a nucleus without also being in contact with a chloroplast (Fig. 2D).

**Figure 2:**
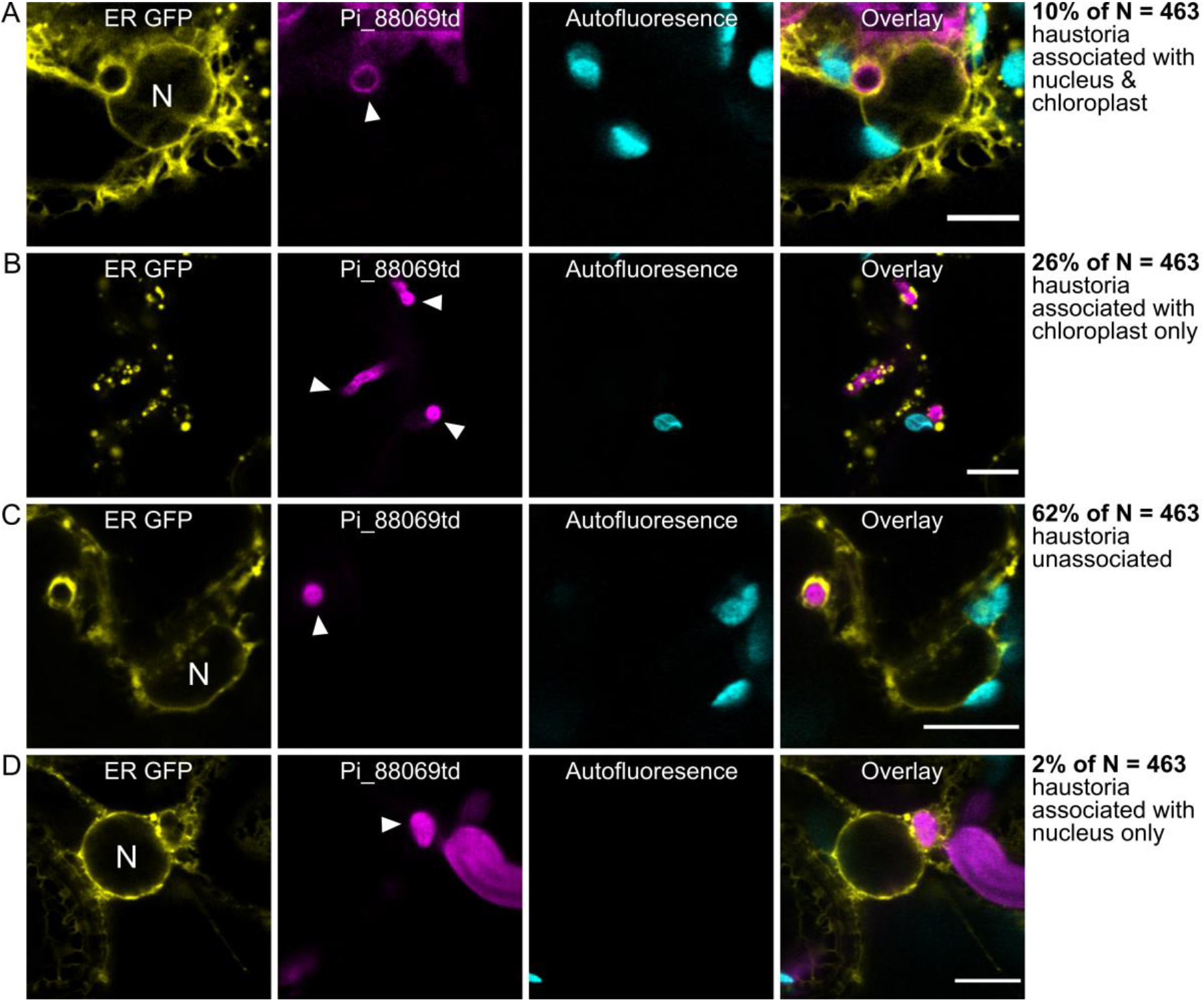
Chloroplasts associate with haustoria both with and without the host cell nucleus during infection. Single plane confocal micrographs of *N. benthamiana* plants expressing GFP in endoplasmic reticulum (ER-GFP), infected with *P. infestans* strain 88069td, showing chloroplast/nucleus positioning around pathogen haustoria. Haustoria marked by white arrow heads, ‘N’ marks the nucleus. Examples of the four combinations of chloroplast/nucleus association with haustoria and the percentage of each observation over 463 haustoria total. Scale bars are 10 μm. **(A)** Dual association of chloroplast and nucleus with haustoria. **(B)** Chloroplast alone at haustoria (no nucleus). **(C)** Haustoria unassociated with either chloroplast or nucleus. **(D)** Nucleus associated with haustoria alone.

Using the same methodology in Figure S1, we compared the frequency of actual chloroplast/nuclear accumulation at haustoria with hypothetical rates expected by chance encounter. We found significantly fewer cases of chloroplast association alone with haustoria than in the mock haustoria data, as well as fewer cases of chloroplast-nucleus co-association with haustoria, but not with nucleus alone association (Fig. S4). Taken together, these results suggest the chloroplast positioning towards haustoria can occur independently of nuclear migration, and that both chloroplast and nuclear accumulation at haustoria occurs at a greater frequency than expected by chance.

### Chloroplasts alter their morphology and contact each other via induction of stromules in response to infection

During live cell imaging of CpGFP plants infected by *P. infestans* 88069td, we also noted an increase in the frequency of stromules – from 4% in the mock infected tissue, to 19% in the infected tissue (Fig. 3A-B). As previously reported, stromules varied in shape and size (M. Schattat et al., 2011); some of these stromules extended towards and wrapped around haustoria (Fig. 3C, Video S9). Further, stromules often extended between different chloroplasts, occasionally even bridging multiple haustoria (Fig. S5, Video S9 & S10). We also monitored stromule-haustorium associations by transiently expressing the EHM marker protein RFP REM1.3 in leaves and infecting with the wild-type *P. infestans* 88069. We noted close association of stromules with the EHM under these conditions (Fig. 3C, Video S11). These results are in agreement with the earlier reports that stromules can be induced by PAMPs (Caplan et al., 2015), and hint at defense-related roles of stromules possibly through mediating chloroplast-chloroplast and chloroplast-EHM associations, as well as the previously reported chloroplast-nucleus associations (Caplan et al., 2015).

**Figure 3:**
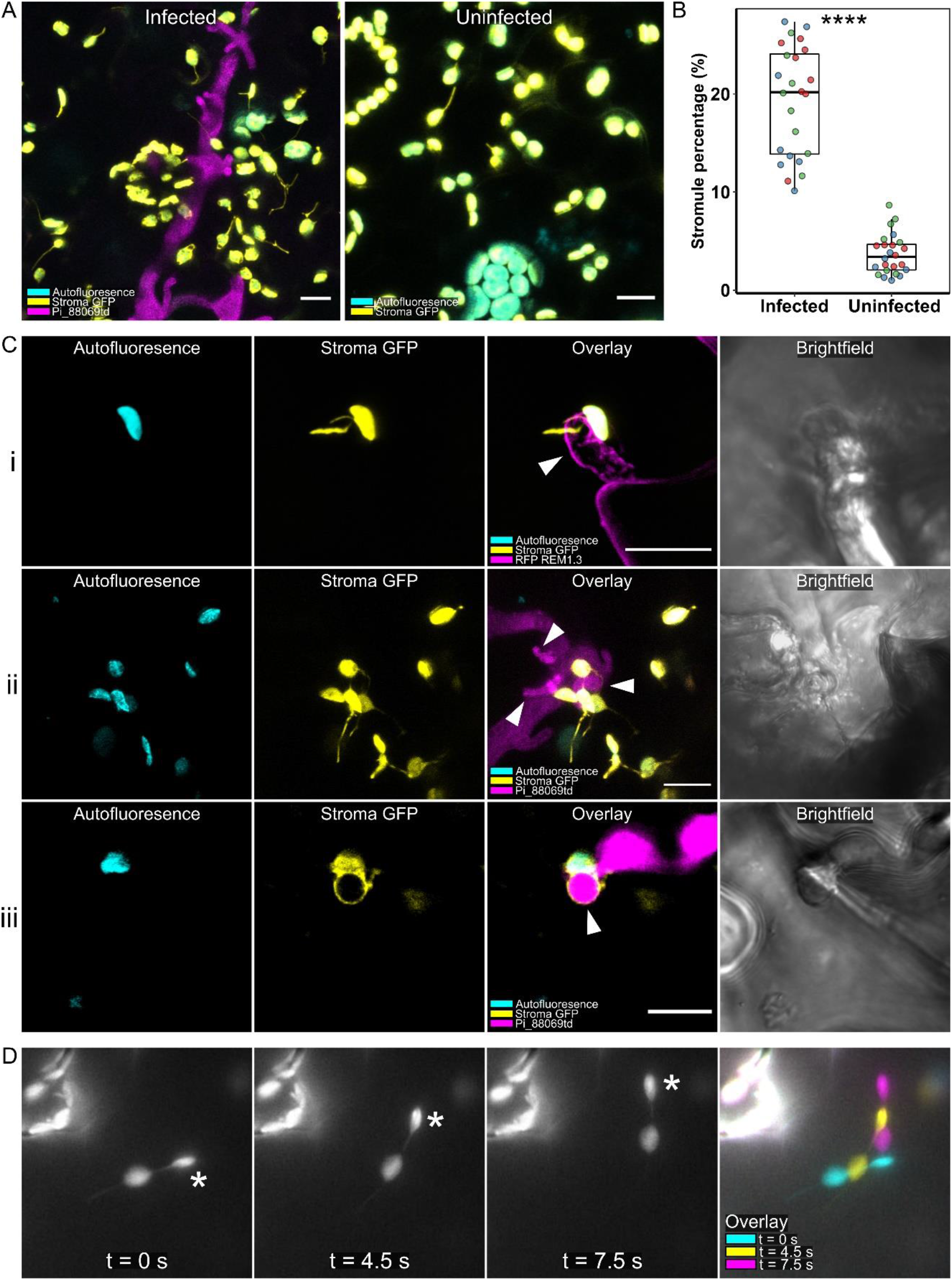
Stromules are induced during infection with *Phytophthora infestans*. **(A)** Maximum projection confocal micrographs of *N. benthamiana* plants expressing GFP in chloroplast stroma (CpGFP), showing induction of stromules during infection with *P. infestans* 88069td. **(B)** Scatter box-plots of the percentage of chloroplasts with stromules for a given image in infected and uninfected samples (exemplified in A). Color of point shows separate biological replicate. Asterisks denote *p* < 0.01 as determined by students t test. **(C)** Maximum projection confocal micrographs of *N. benthamiana* plants expressing GFP in chloroplast stroma (CpGFP), infected with *P. infestans*, showing examples of stromules wrapping around haustoria. Haustoria marked by white arrow heads. i. Infection with wild-type *P. infestans* 88069, plant cell transiently expressing RFP REM1.3 to mark the extrahaustorial membrane. ii-iii. Infection with *P. infestans* 88069td. **(D)** GFP channel in grayscale from TIRF microscope. Laser capture of chloroplast (asterisk) linked by stromule to another chloroplast. When the trapped chloroplast moves, the linked chloroplast co-migrates. All scale bars are 10 μm.

During our attempts to move chloroplasts in infected cells, we once attempted to move a chloroplast that was seemingly connected to another by a stromule like extension. Here, we observed co-migration of the chloroplasts by moving only one of the pair (Fig. 3D, Video S12), indicating that chloroplasts may be linked by their stromules, or that chloroplasts can follow one another when bridged by a stromule. Co-migration between non-stromule connected chloroplasts has also been previously observed (Caplan 2018). However whether plastids can fuse to form a continuous stromal compartment that enables macromolecule exchange thorough stromules is still under debate (Hanson & Hines, 2018; M. H. Schattat et al., 2015).

### Stromules are induced upon surface immune activation

PAMPs trigger a range of immune responses when recognized by surface localized immune receptors (Jones & Dangl, 2006). Flg22, a peptide PAMP from bacterial flagellin, was previously shown to induce stromules (Caplan et al., 2015). We replicated this result with flg22, while also testing other elicitors: chitin, a polysaccharide PAMP of fungal microbes and arthropod pests; and INF1, an extracellular *P. infestans* protein that, unlike most PAMPs, elicits host cell death upon perception by the plant (Kamoun et al., 1998). Flg22, chitin, and INF1 all induced stromules after 24 hr compared to the water control (Fig. 4A), suggesting stromule production is a general response to a range of microbes. PAMP activity was confirmed by detection of phosphorylated mitogen-activated kinases (MAPKs) by Western blot (Fig. S6).

**Figure 4:**
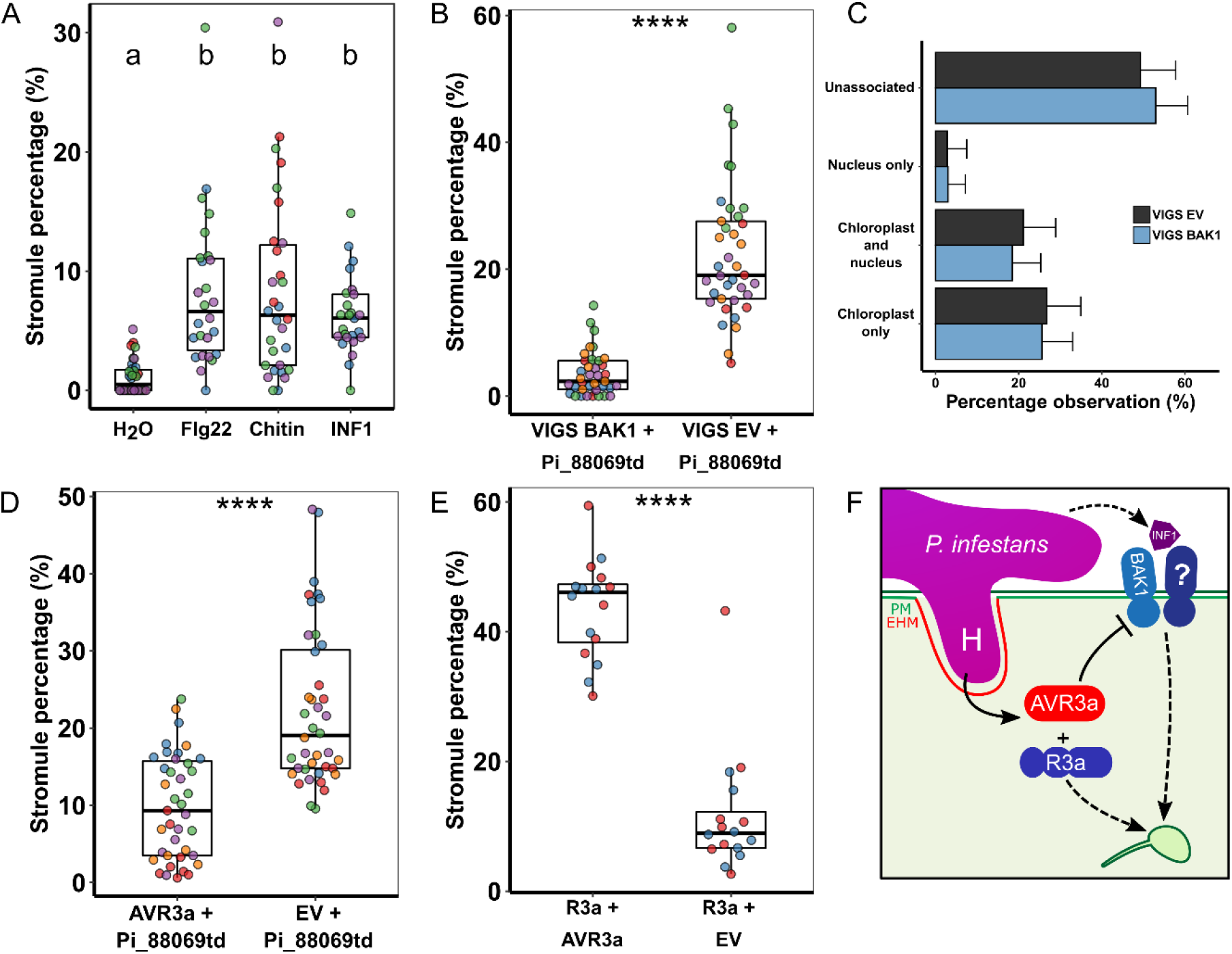
Stromule induction through immune recognition. **(A)** Scatter box-plots of the percentage of chloroplasts with stromules for a given image following 24 hr treatment of CpGFP plants with water, flg22, chitin, or INF1. Color of point shows separate biological replicate. Letters mark significance groups (where α = 0.01) as determined by Dunn’s test with p-values adjusted by Bonferroni correction. **(B)** Scatter box-plots of the percentage of chloroplasts with stromules for a given image following during infection of CpGFP plants by *P. infestans* strain 88069td in VIGS *BAK1*, and VIGS EV (control) plants. Color of point shows separate biological replicate. Asterisks denote *p* < 0.01 as determined by Wilcoxon test. **(C)** Bar plots showing percentage of chloroplast/nuclear accumulation at haustoria in VIGS *BAK1* plants, compared to VIGS EV plants. Observations made in 16C-GFP plants infected with *P.* infestans strain 88069td, across 3 separate biological replicates, totaling 168 and 142 haustoria in the VIGS *BAK1* and VIGS EV conditions respectively. Error bars show confidence intervals. No statistically significant difference detected (α = 0.05), as determined by Fisher’s Exact test. **(D)** Scatter box-plots of the percentage of chloroplasts with stromules for a given image following during infection of CpGFP plants by *P. infestans* strain 88069td during transient plant cell expression of either AVR3a or the EV control. Color of point shows separate biological replicate. Asterisks denote *p* < 0.01 as determined by Wilcoxon test. **(E)** Scatter box-plots of the percentage of chloroplasts with stromules for a given image following transient co-expression of either R3a with AVR3a, or R3a with an EV control. Color of point shows separate biological replicate. Asterisks denote *p* < 0.01 as determined by Wilcoxon test. **(F)** Model of possible immune related signaling pathways that can lead to stromule induction, evidenced by data shown in A, B, D & E.

BAK1 is a surface localized co-receptor that mediates immune signaling in co-operation with various pattern-recognition receptors to perceive PAMPs (Chaparro-Garcia et al., 2011; Heese et al., 2007). Therefore, we monitored stromule formation upon systemic silencing of *BAK1* by virus induced gene silencing (VIGS) in CpGFP plants (Fig. S7) during infection (Fig. 4B). We measured a substantial decrease in infection triggered stromule induction following *BAK1* silencing (4%, *N* = 37 images quantified) compared to control silencing (23%, *N* = 37 images quantified), corroborating the induction of stromules by PAMPs and suggesting that PAMPs from *P. infestans* induce stromules during infection.

Extending this, we tested whether silencing of *BAK1* through VIGS would alter chloroplast and nucleus accumulation at the haustorium. To do this, we used live cell infection microscopy of *BAK1* silenced ER-GFP plants. We found no significant difference in chloroplast and/or nuclear accumulation at haustoria during infection compared to silencing controls (*N* = 168 and *N* = 142 haustoria respectively) (Fig. 4C). This indicates that BAK1 dependent signaling for stromule induction is independent from that of chloroplast accumulation at the host-pathogen interface.

To complement the *BAK1* silencing phenotypes on stromule formation, we then tested whether AVR3a, a host-translocated effector of *P. infestans* that suppresses BAK1-mediated immune signaling (Chaparro-Garcia et al., 2011, 2015), can perturb pathogen induced stromule development. Following transient expression of AVR3a, stromule formation during infection decreased significantly, from 23% to 10% (Fig. 4D). This indicates pathogen effectors can perturb stromule induction, further supporting a defense-related role of stromules.

Functionality of AVR3a was confirmed by observing cell death when transiently co-expressed with the cognate R3a NLR immune receptor (Fig. S8). We also observed that in the absence of the pathogen, transient co-expression of AVR3a with R3a led to an induction of stromules compared to the control, where R3a was transiently co-expressed with EV instead of AVR3a (Fig. 4E). This points towards the induction of stromules by HR cell death triggered by effector recognition, despite the functionality of the effector in suppressing stromules when it goes undetected by the plant. Taken together, these results show importance of cell-surface immune perception of PAMPs in the induction of stromules and suggests this process can be suppressed by pathogen effectors but can be elicited upon the detection of those effectors (Fig. 4F).

### HyPer ROS reporter displays increased reactivity in chloroplasts upon infection

Previously, Caplan et. al. (2015) proposed that close proximity of chloroplasts to the plant nucleus could allow for ROS transfer to the nucleus via stromules. We reasoned that chloroplast positioning around the haustorium may serve to increase localized reactive oxygen species (ROS) production by chloroplasts at the site of infection. To test this, we visualized ROS in live cell infection using the HyPer ROS sensor fused to the chloroplast transit peptide of *A. thaliana* RecA, cTP-HyPer herein (Caplan et al., 2015). In the presence of ROS, the ratio between emission intensity from 405 nm and 488 nm excitation changes, giving a detectable readout of ROS from confocal microscopy (Belousov et al., 2006).

The epidermal plastids of infected cells showed noticeably more signal than the uninfected cells (Fig. 5). However, within a single haustoriated cell, we could not see a discernable difference in ROS signal between the different plastids, including between those associated and those unassociated with haustoria (Fig. 5). We speculate that chloroplast ROS production and visualization may be highly variable due to differences in stages of infection and the changes in the microenvironment as well as potential manipulation by effector secreted by the pathogen.

**Figure 5:**
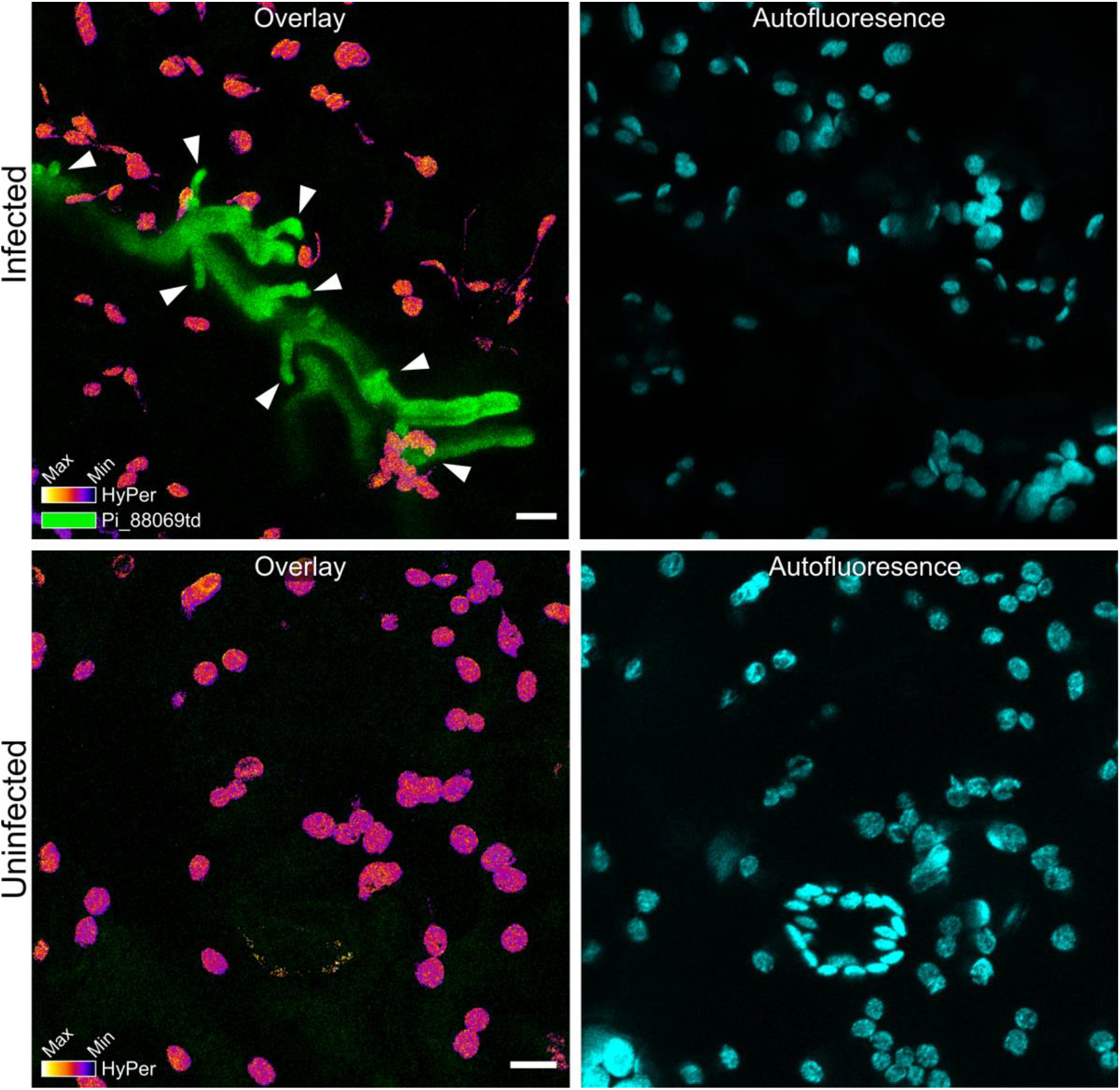
Chloroplasts increase ROS production during infection. Maximum projection confocal micrographs of wild-type *N. benthamiana* plants transiently expressing the plastid localized cTP-HyPer ROS sensor, with and without live infection by *P. infestans* strain 88069td. Visual representation of the cTP-HyPer signal intensity ratio between excitation by 405 and 488 nm lasers, shown with the ImageJ ‘Fire’ LUT. Haustoria marked by white arrow heads. All scale bars are 10 μm. All confocal settings were kept identical between infected and uninfected during acquisition and all channels were displayed, hence why some green background signal is visible in the uninfected image.

## Discussion

Here, we demonstrate that during host colonization by *P. infestans*, chloroplasts accumulate at the pathogen interface (Fig. 1-2) and alter their morphology through induction of stromules (Fig. 3-4). Additionally, we show that nuclei are almost exclusively localized to haustoria in the company of a chloroplast, but that chloroplasts can accumulate independently (Fig. 2). Stromules occasionally embrace the EHM and link chloroplasts to each other, forming dynamic organelle clusters around the pathogen interface (Fig. 3C, Fig. S5) that are reminiscent of mitochondrial networks with poorly understood functions (Hoitzing et al., 2015). Notably, infection-triggered stromule development relies on surface immune signaling, whereas the pathogen can subvert this process remotely interfering with these pathways (Fig. 4). Our results implicate chloroplasts in cell polarization upon pathogen attack and point to more complex functions of these organelles in plant-pathogen interactions.

### How do chloroplasts position at the pathogen interface?

It is well-established that chloroplasts alter their subcellular localization in response to light in an actin dependent manner (Kadota et al., 2009; Wada & Kong, 2018). We found that infection induced chloroplast positioning around the haustorium relies to an extent on actin polymerization (Fig. 1B-C), whereas the process is not affected by light conditions (Fig. S3). Previous studies have shown that modulation of the actin cytoskeleton can affect cell penetration by filamentous plant pathogens (Kobayashi & Hakuno, 2003; Tang et al., 2016), and others have noted the re-organization of actin around the site of pathogen penetration (Opalski et al., 2005). Our finding, that actin polymerization inhibitor LatA reduces chloroplast positioning at haustoria (Fig. 1C) is in agreement with previous studies that implicate actin in the focal immune response, as well as the known importance of actin in both chloroplast movement (Higa et al., 2014; Suetsugu et al., 2016; Suetsugu & Wada, 2016). While this is a promising first step in altering chloroplast positioning during infection, we also heed caution in the use of pharmacological treatment of live infection systems with potential off target effects in both host and pathogen. Ideally, work following up to this would move towards using targeted genetic approaches or identifying effectors that can target this process more specifically.

Additionally, we monitored chloroplast association to haustoria in the context of nuclear association (Fig. 2). Although the movement of nuclei towards plant-pathogen interfaces appear to be under varying spatio-temporal dynamics in different patho-systems (Griffis et al., 2014; Scheler et al., 2016), evidence suggest that this process may contribute to plant immunity (Daniel & Guest, 2006). However, nuclear movement during cell penetration by filamentous microbes is not exclusively uni-directional, with the nucleus moving first towards and then away from penetration sites in many interactions (Genre et al., 2005; Schmelzer, 2002). Our results regarding the accumulation of nuclei provide a snapshot at a single time point of infection, therefore our conclusions are derived from the observed state of the nucleus at that time point, not considering whether the nucleus is in the process of moving towards or away from the haustorium. Further dissection of specific nuclear movement components over extended time courses is required to address the intricacies and impact of nuclear movement in plant focal immunity.

### Why do chloroplast alter their morphology during infection?

Our findings on PAMP induction of stromules (Fig. 4A) validate and expand upon those from Caplan et. al. (2015), who previously showed that flg22 can induce stromules. The reduction of stromules during infection by systemic silencing of *BAK1* (Fig. 4B) suggest that a major sub population of stromules (if not all) induced during infection rely on BAK1 mediated immune signaling initiated at the cell surface. Here, we used AVR3a, an effector protein, as a tool to cross-examine the role of BAK1 (Fig. 4B) since it is known to suppress BAK1-mediated defense-signaling (Chaparro-Garcia et al., 2011, 2015). Furthermore, we hypothesize that other effector proteins are likely able to inhibit stromules indirectly by targeting similar cell surface signaling pathways.

Interestingly, *BAK1* silencing did not affect positioning of chloroplasts and/or the nucleus around haustoria (Fig. 4C), hinting at differential signaling pathways between stromules and organelle positioning. As of yet, we do not know what signaling takes place to re-route plant defenses towards the host-pathogen interface. The decoupling of stromule induction and chloroplast accumulation at haustoria is further supported by our time-lapse imaging which shows accumulation of chloroplasts at haustoria can occur independently of stromules, although we observe both stromule led and stromule independent movement towards haustoria (Video S3 & S4). This is in keeping with observations by Kumar et. al. (2018), who found that chloroplast movement is mostly, but not exclusively, stromule directed.

Induction of stromules by immune signaling during pathogen attack strongly points to defense-related functions of these tubular organelle extensions. But how could stromules contribute to immunity? While our understanding of stromules is still limited by a lack of stromule specific inhibitors/inducers, a model of stromules as signaling conduits is emerging. Caplan et. al. (2015) suggest that the increased surface area provided by a stromule aids in the transfer of chloroplast synthesized pro-defense molecules to the cytosol and nucleus where they function. The increase in chloroplast-nucleus contact, facilitated by stromules, and triggered during an HR response, is thought to amplify the progression of HR in a positive feedback loop. Similarly a more recent study tracked the spatiotemporal redox state of chloroplasts, and stromule induction, in potato during potato virus Y challenge (Lukan et al., 2021). Lukan et. al. (2021) conclude that, in this patho-system, stromules are involved in signaling on the virus multiplication front.

We observed that chloroplasts establish network-like interactions via stromules (Fig. S5), and that some chloroplasts intimately associate with the haustorium interface through stromules (Fig. 3C). These chloroplast clusters and stromule extensions around the haustorium could plausibly aid the coordination of defense-related functions of chloroplasts by, for instance, mediating deployment of pro-defense molecules at the pathogen interface. Our findings further support the notion that stromules are induced to contribute to pathogen defense (Caplan et al., 2015; Erickson et al., 2018), but the function of stromules still remains to be determined (Hanson & Hines, 2018).

### Right time, right place: Chloroplast position at pathogen interface is a host defense or pathogen strategy?

It is unclear whether the association of chloroplasts with the haustoria of *P. infestans* is a plant defense or a pathogen virulence strategy given the arsenal of immune chemicals produced by chloroplasts, it is plausible that their presence at haustoria may enhance the effectiveness of their deployment. However, we cannot discount the possibility that chloroplasts accumulate at haustoria to the benefit of the pathogen, perhaps serving to nourish the parasite. Specific, genetic strategies that impair chloroplast positioning around the haustorium are necessary to reach definitive conclusions on this. Expanding this work to other patho-systems, particularly those distantly related or those with a different lifestyle (i.e. completely biotrophic), as well as using alternative strains of *P. infestans* with different effector repertoires would also help resolve the physiological and evolutionary relevance of this observed response. We believe this work will lay the foundation for future studies regarding chloroplast movement towards and association with intracellular pathogen structures. By further defining the role of surface immune signaling in stromule induction and showing how effectors can be used to manipulate this process, we believe these tools will help accelerate research into the stromule function and signaling pathways.

## Materials and Methods

### Biological Material

*Nicotiana benthamiana* plants grown in a growth chamber at 25°C under high light intensity (16-h-day/8-h-dark photoperiod). Transplastomic GFP-expressing *N. benthamiana* plants, accumulating GFP in the chloroplast stroma (Stegemann et al., 2012), and transgenic GFP-expressing *N. benthamiana* plants, accumulating GFP in the endoplasmic reticulum were maintained in the same conditions as wild-type *N. benthamiana. Phytophthora infestans* isolate 88069 (Van West et al., 1998) and 88069td (Whisson et al., 2007), a transgenic strain expressing the red fluorescent marker tandem dimer RFP (tdTomato), were used. Both isolates were cultured on plates with rye sucrose agar (RSA) for 12-16 days at 18°C in the dark, as described elsewhere (Song et al., 2009) prior to use for infection of *N. benthamiana*.

### Plasmid constructs

The following constructs used in this study were previously published as follows: RFP:REM1.3 (Bozkurt et al., 2014); R3a (Chaparro-Garcia et al., 2015); AVR3a cloned in pICSL86977 was provided by TSLSynBio; GFP Actin chromobody (Rocchetti et al., 2014). Silencing construct TRV2-BAK1 was kindly provided by The Sainsbury Lab (Chaparro-Garcia et al., 2011). RecA-cTP HyPer construct was kindly provided by Prof. Savithramma Dinesh-Kumar (Caplan et al., 2015).

### Transient gene-expression assays in *N. benthamiana*

*Agrobacterium tumefaciens* GV3101 strain (Hellens et al., 2000) carrying T-DNA constructs was used to mediate transient gene expression (referred to in text as transient expression) into 3-4-week-old *N. benthamiana* leaves, as previously described (Bozkurt et al., 2011, 2014). Briefly, overnight cultures of transformed *A. tumefaciens* were washed and harvested with 1500 μL autoclaved dH_2_O by centrifugation at 1500 *g* twice and resuspended in agroinfiltration buffer (10 mM 2-(*N*-morpholino-ethanesulfonic acid hydrate (MES hydrate), 10 mM MgCl_2_, pH 5.7). For the transient expression assays, each *A. tumefaciens* construct was mixed in agroinfiltration buffer to achieve a desired final OD_600_ each *A. tumefaciens*. For GFP Actin, OD_600_ = 0.05; for AVR3a, R3a, and EV, OD_600_ = 0.3. For RFP REM 1.2, OD_600_ = 0.3. For cTP-HyPer, OD_600_ = 0.2. *P. infestans* inoculations were performed 4 to 24h after infiltrations if at all.

### Virus induced gene silencing (VIGS)

*Agrobacterium* was prepared as above carrying TRV1 and the appropriate TRV2 construct and mixed to a final OD_600_ of 0.4 or 0.2 respectively, in agroinfiltration buffer supplemented with 100 μM acetosyringone (Sigma) and left in the dark for 2 h prior to infiltration to stimulate virulence. 14-day old *N. benthamiana* seedlings were infiltrated in both cotyledons and any true leaves that had emerged. *N. benthamiana* plants were infiltrated with TRV1 and TRV2-BAK1 for *BAK1*-silencing and TRV1 and TRV2-EV for the empty vector control. TRV2 containing the *N. benthamiana* sulfur (Su) gene fragment (TRV2-NbSU) was used as a positive control to indicate viral spread. Plants were left to grow under standard conditions until experiments could be carried out four weeks later.

### *Phytophthora infestans* infection

Zoospores were harvested from sporangia by addition of cold distilled water and collected after 2h of incubation at 4°C, adjusting dilution to 50,000 spores/ml. Infections were performed by the addition of 10 μL of zoospore droplets to the abaxial side of the leaf. The infected leaves were maintained in plastic boxes on damp paper towels at 18°C under 16-h-day/8-h-night conditions (except in the dark/dark experimental condition where there was no day period, Fig. S3).

### PAMP treatments

Flg22 and INF1 purified peptides were provided by The Sainsbury Laboratory (Norwich). Chitin was prepared from powdered shrimp shell (Sigma-Aldrich) as per manufacturers protocol. Working concentrations of 1 μM (flg22 and INF1) and 100 μg mL-1 (chitin) were used unless otherwise stated. Dilutions were made in water and infiltrated into the underside of leaves using a needless syringe.

### LatA treatment

Latrunculin A (abcam) was diluted to a stock concentration of 100 μM in 100 % DMSO. Water control was prepared with matching final concentration of DMSO (v/v). LatA was infiltrated into leaf tissue by needless syringe, 24 hours before microscopy.

### Visualization of chloroplast ROS

Live cell imaging of chloroplast ROS was imaged using the HyPer ROS sensor (Belousov et al., 2006) fused to the chloroplast transit peptide of *Arabidopsis thaliana* RecA. This ROS sensitive fluorescence-based marker is imaged by fast line switching between 405 nm (channel 1) and 488 nm excitation (channel 2), detecting emission in the range of 491–543 nm. The ratio between emission from channel 1 and channel 2 gives the final signal by dividing signa from channel 2 by channel 1 in ImageJ using the inbuilt ‘Math’ functions (Schneider et al., 2012). Lookup table was set to Fire in ImageJ for better visualization of intensity.

### RT-PCR assay

60 mg of leaf tissue was excised from 5-week old leaves (VIGS experiments) and frozen in liquid N2. RNA was extracted from the leaf tissue using the Plant RNA Isolation Mini Kit Protocol (Agilent Technologies). RNA quality and concentration was measured using a NanoDropTM Lite Spectrophotometer (Thermo Scientific). cDNA was synthesized using as a template 2 μg of RNA following the SuperScript II RT protocol (Invitrogen). To amplify the cDNA, a standard PCR (RT-PCR) was then performed using DreamTaq DNA polymerase (5 U/μL) (Thermo Scientific). VIGS *BAK1* silencing was confirmed as previously described (Chaparro-Garcia et al., 2011).

### Confocal microscopy

All microscopy analyses were performed on live *N. benthamiana* epidermal cells 2-6 days post agroinfiltrations and infections. Leaf discs were excised and imaged on either a Leica SP5 or SP8 resonant inverted confocal microscope (Leica Microsystems) using 63X, or 40X respectively, 1.2NA Plan-Apochromat water immersion objective. Specific excitation wavelengths and filters for emission spectra were set as described previously (Koh et al., 2005). The Argon laser excitation was set to 488 nm and the Helium-Neon laser to 543 nm and their fluorescent emissions detected at 495–550 and 570–620 nm to visualize GFP and RFP fluorescence, respectively. To avoid bleed-through from different fluorophores, images were acquired using sequential scanning and Maximum Intensity Projections were created from the Z-stacks. 3D images and videos were generated with confocal files in 12-bit TIFF format imported into NIS-Elements (Version 4.50, Nikon Instruments, UK) and processed with Advanced Denoising. Videos were made using the Volume View and Video Maker modules.

### Optical trapping setup

Optical trap for chloroplast/stromule capture was setup as described by Sparkes et al 2017, Chapter 13 (Sparkes et al., 2018). An optical trap with a two-channel TIRF microscope (TIRF-M) was combined with a Nikon Ti-U inverted microscope. Optical trapping was performed using a near infrared trapping laser at 1070 nm using a Nikon 100x, oil immersion, NA 1.49 TIRF objective lens. For GFP and RFP chromophores fused to the proteins of interest were excited using 488 and 561 nm laser diode, respectively. Their Fluorescent emissions were detected using two electron multiplying charge-coupled device (EMCCD, iXon, Andor) cameras. The sample (~5 mm^2^ leaf tissue) was mounted on a computer-controlled variable speed (Märzhäuser) stepper motor stage. The associated computer-controlled hardware was interfaced using National Instruments LabVIEW which provides full automation for each trapping routine. The power of the optical trap laser transmission was set to 40.7 mW. The TIRF image was recorded from 0 s, the trap was turned on at 1 s, the translation stage movement of 10 μm at 2 μm/s begins at 5 s and ends at 10 s, the trap was deactivated at 11 s, and the image recording stops at 22 s (relating to 11 s recovery periods). A 10 μm distance threshold was chosen to ensure consistence between experiments by which chloroplasts were moved; in particular, 10 μm was chosen as it is not so far as to generate potential side effects such as pushing the chloroplast into the tonoplast, yet large enough to move chloroplasts a substantial distance from their original position.

### Micrograph quantification

Chloroplast stromule quantification was done automatically using a MATLAB script. Stromules were manually counted using a semi-automated MATLAB script. Percentage of chloroplasts with stromules were calculated by dividing the number of chloroplasts one (or more) stromule(s) by the total number of chloroplasts.

Quantification of haustorial-chloroplast-nucleus accumulation was done manually from original confocal micrographs by looking through each layer of the z-stack with only the brightfield and *P. infestans* 88069td channels active to identify haustoria without bias. Association of chloroplasts and or nuclei to these marked haustoria was then counted, assessing each layer of the z-stack as opposed to viewing a z-projection that could cause false-positives. For Fig. 1C & S3B, two individuals independently quantified chloroplast-haustoria association, with discrepancies re-checked to reach a consensus quantification.

### Mock haustoria

Mock haustoria were applied to each of the confocal micrographs that were used in the data sets that contributed to Fig. 1A and Fig. 2 using standard ImageJ line tools. All channels were turned off except the brightfield and the *P. infestans* 88069td channel to reduce bias. To position the mock haustoria, a straight line was drawn that bisected the base (where the haustoria enters the plant cell) to the tip of each actual haustoria, extending across the vacuole until the cell border opposite was hit (as seen from the brightfield). This end position was taken to be the position at which a mock haustoria penetrated the cell. The line width was set to 3 μm (based on the general observation that haustoria were 2.7 μm in width). All channels were turned back on and the region around the mock haustoria and the instances of chloroplast-nucleus presence adjacent to these mock haustoria were counted, note only the z-slices containing the actual haustoria from which the mock was positioned were counted.

This method was used to keep the number of mock and actual haustoria per cell similar for comparison, but in the following cases, a mock haustoria could not be placed: if the mock haustoria position overlaps with an actual haustoria, if the cell border in out of the field of view, if the actual haustoria has developed in the crook of a cell and is touching both cell borders. If no mock haustoria could be successfully placed, the image was not included in the pair-wise comparison.

### Statistical analysis

Statistical significance of the differences observed when comparing means (stromule quantification) were assessed by t test when found to be normal by Shapiro-Wilk test. If data was found to be non-normally distributed Wilcox statistical test was implemented by R. A pair-wise Wilcoxon test was used to compare mock v actual haustoria accumulation (Fig. S1B, Fig. S2), with samples paired by data coming from the same micrograph. Counts of haustoria-chloroplast-nucleus association were pooled together to generate an overall proportion/percentage from all micrographs instead of treating each micrograph as technical repeat and its taking percentage association as a single data point; this was done to avoid data skew by micrographs that contained only one haustoria and would therefore generate many 100% and 0% values that skew the mean estimate. The proportions of each observation were compared using a Fisher’s Exact test in R where statistical comparison was made.

### Chloroplast automated counting algorithm through image processing

The image processing algorithms were used to calculate the gradient of the image to identify the boundaries of the puncta. Enclosed regions formed by the boundaries were algorithmically identified and counted. This procedure was done for each individual channel green (in chloroplast stroma) and blue (Chloroplast Auto-fluorescence). The chloroplast (GFP channel) containing stromule were counted in a semi-automated fashion.

### Western blotting

Protein extraction, purification and western blot analysis steps were performed as described previously (Bozkurt et al., 2011). Anti-phosphorylated MAPK (Phospho-p44/42 MAPK, Cell Signaling Technology) was used as primary antibody, anti-rabbit (Sigma-Aldrich, UK) antibody was used as secondary.

## Acknowledgements

We thank Dr. Alex Jones (Warwick) for initiating collaboration with IS, Dr. Sebastian Schornack (SLCU) for initiating collaboration with MS, Prof. Peter Nixon (Imperial) for providing CpGFP plant seeds, Prof. Savithramma Dinesh-Kumar for providing us with the RecA-cTP HyPer construct.

## Funding

Bozkurt lab funded by BBSRC (BB/M002462/1). The Facility for Imaging by Light Microscopy (FILM) at Imperial College London is part-supported by funding from the Wellcome Trust (grant 104931/Z/14/Z) and BBSRC (grant BB/L015129/1).

## Author contributions

Conceptualization: ZS, CD, MES, MS, IS, TOB; Data Curation: ZS, CD, AT, PP, LHY, DCAG, BCB, IS, TOB; Formal Analysis: ZS, CD, AT, PP, YL, BCB, IS, TOB; Funding acquisition: TOB; Investigation: ZS, CD, AT, PP, YL, MES, LHY, AYL, VK, BCB, TOB; Methodology: ZS, CD, AT, YL, DCAG, ADW, SWB, BCB, MS, IS, TOB; Project Administration: ZS, CD, TOB; Resources: ZS, CD, AT, AYL, VK, IP, IS, TOB; Software: ZS, IP; Supervision: ZS, CD, AT, PP, MES, BCB, IS, TOB; Validation: ZS, CD, AT, PP, YL, MES, YT, ADW, MS, IS; Visualization: ZS, CD, TOB; Original Draft Preparation: ZS,CD, TOB; Review & Editing: ZS, CD, AT, PP, MES, LHY, DCAG, AYL, ADW, SWB, MS, IS, TOB.

## Competing interests

Authors declare no competing interests.

## Data and materials availability

All data is available in the main text, the supporting information or other raw data is available upon request. We are happy to provide all materials used here upon request.

## Supplementary information

**Figure S1:**
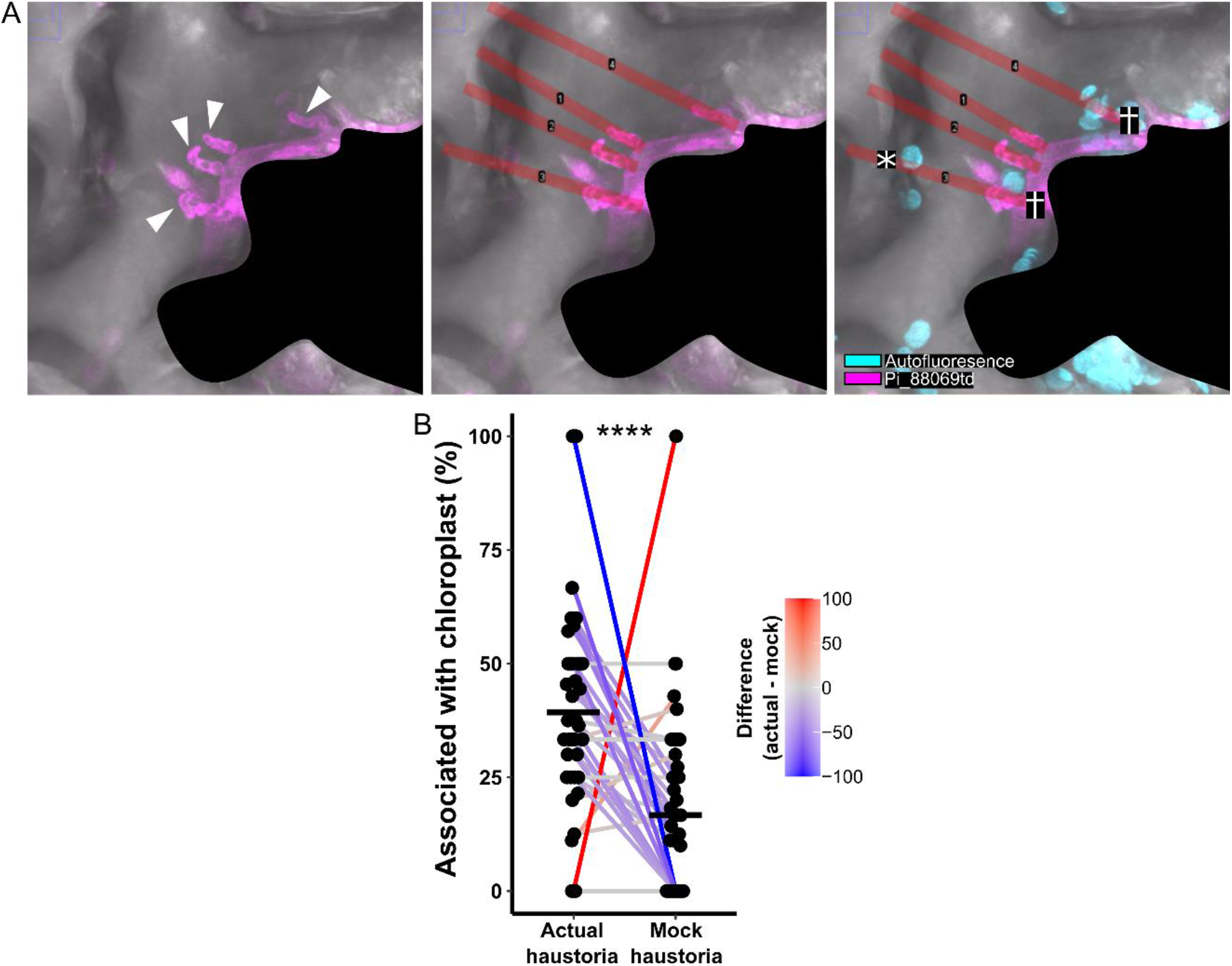
Mock haustoria positioned opposite real haustoria are less frequently adjacent to chloroplasts. **(A)** Confocal micrograph of *N. benthamiana* infected with *P. infestans* strain 88069td as an example of how mock haustoria were positioned and used to assess the rates of chloroplast-haustoria association that might be expected by chance encounter. White arrow heads show actual haustoria in the cell. Red lines show trajectory from an actual haustoria to the opposite site of the infected cell, the point at which a mock haustoria was then marked. Cross (†) shows association of actual haustoria with a chloroplast, asterisk (*) shows the association of a mock haustoria with a chloroplast. Shaded area is another infected cell, removed to simplify the explanation of the method. **(B)** Paired samples scatter plot showing the image-by-image percentage of chloroplast association between the actual and mock haustoria (*N* = 39 images total). Black cross bar marks the mean. Samples are paired based on derivation from the same confocal image, paired samples are joined by a line that is colored based on the value difference between them. Asterisks denote *p* < 0.01 as determined by a paired samples Wilcoxon test.

**Figure S2:**
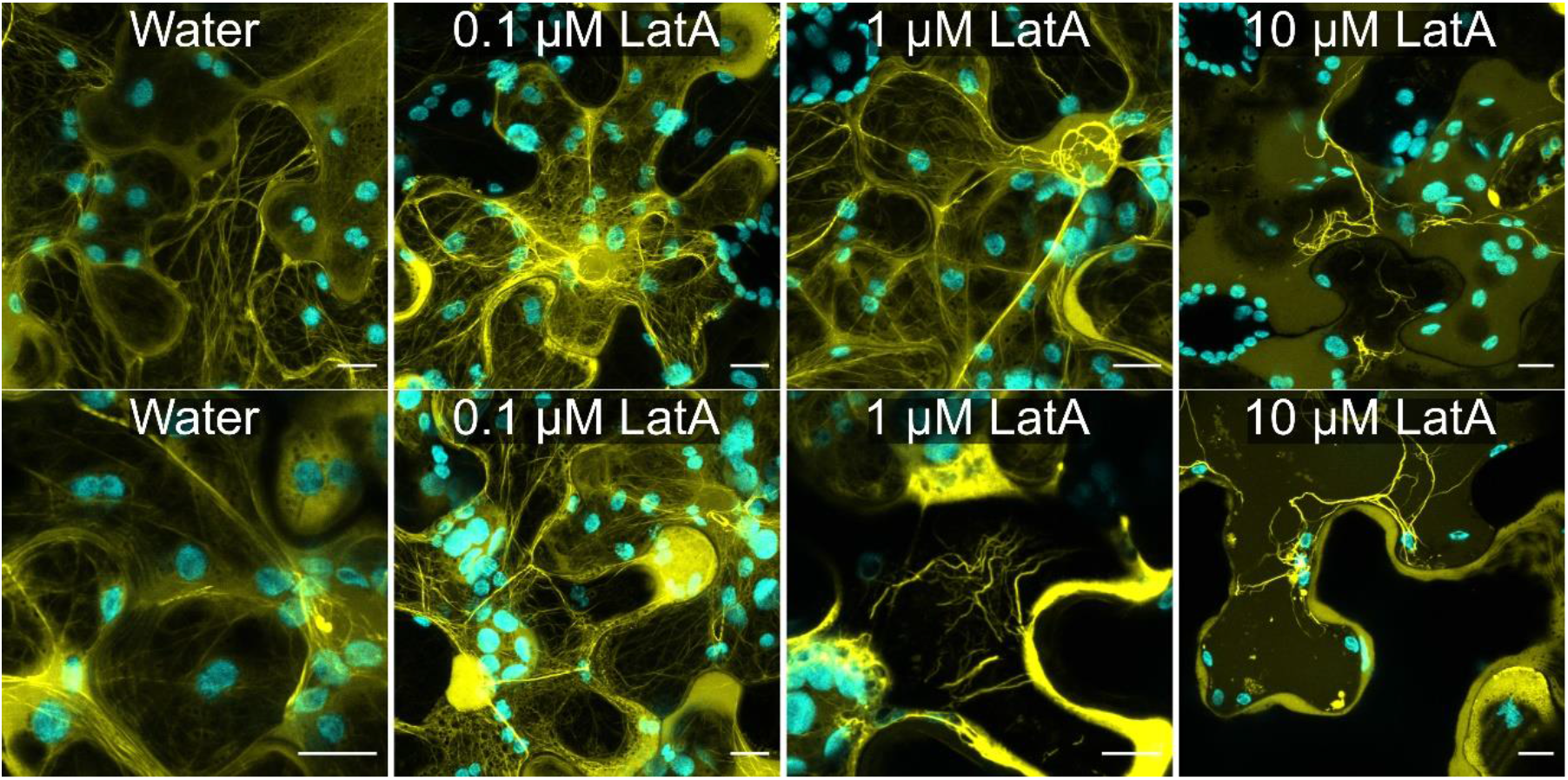
Increasing concentrations of LatA cause increasingly visible disruption to the normal actin filaments. Maximum projection confocal micrographs of wild-type *N. benthamiana* plants, transiently expressing GFP actin chromobody, following 24 hr treatment with water, 0.1, 1 or 10 μM LatA. Scale bars are 10 μm. Two example images shown for each condition. Cyan shows chlorophyll autofluorescence, yellow shows GFP actin. All scale bars are 10 μm.

**Figure S3:**
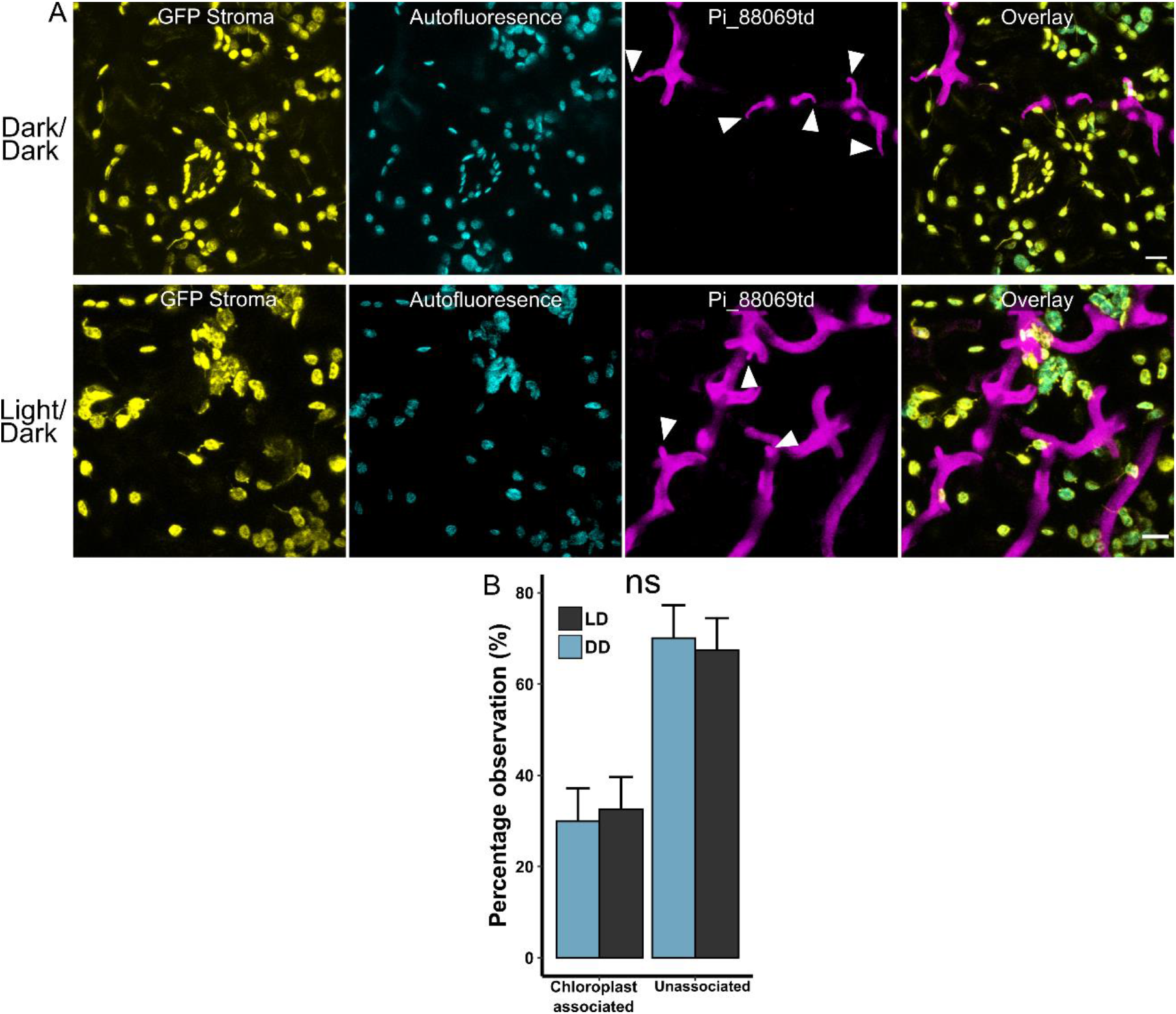
Alternative lighting regimes do not affect the association of chloroplasts with haustoria. **(A)** Maximum projection confocal micrographs of *N. benthamiana* plants expressing GFP in chloroplast stroma (CpGFP), infected with *P. infestans* strain 88069td, following differential lighting regimes. Either, two days in the dark (dark/dark) or normal day/night cycling (light/dark) for the same amount of time. Haustoria marked by white arrow heads. Scale bars are 10 μm. **(B)** Bar plots showing percentage of chloroplast accumulation at in different lighting regimes. Observations made in *N. benthamiana* plants expressing GFP in chloroplast stroma (CpGFP) infected with *P.* infestans strain 88069td, across 2 separate biological replicates, totaling 175 and 167 haustoria in the light/dark and dark/dark conditions respectively. Error bars show confidence intervals. No statistically significant difference detected (α = 0.05), as determined by Fisher’s Exact test.

**Figure S4:**
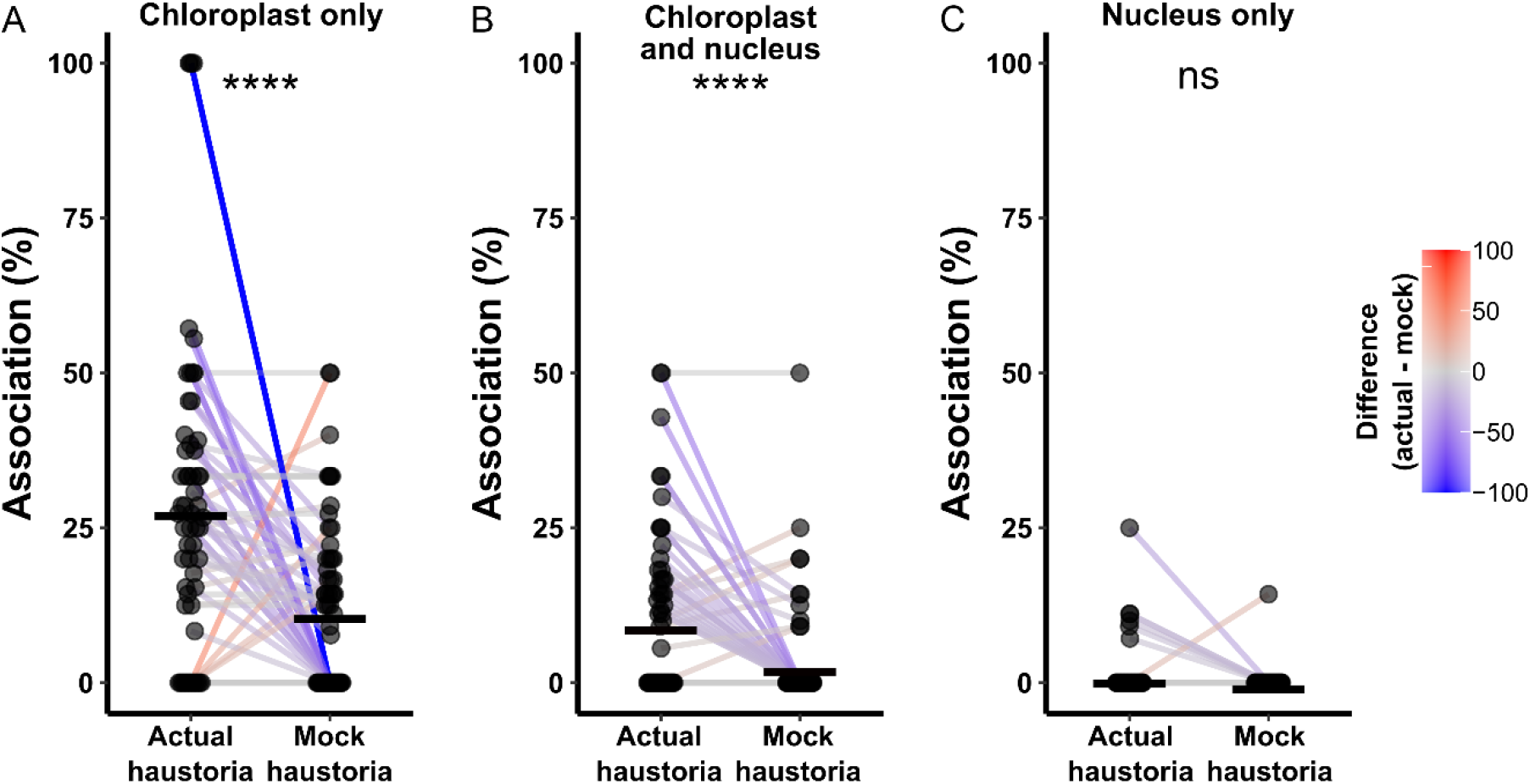
Haustoria association with chloroplasts alone, or chloroplasts and the nucleus together, occurs at a higher frequency than expected from chance. Paired samples scatter plot showing the image-by-image percentage of haustoria association between the actual and mock haustoria (*N* = 60 images total). Black cross bar marks the mean. Samples are paired based on derivation from the same confocal image, paired samples are joined by a line that is colored based on the value difference between them. Asterisks denote *p* < 0.01 as determined by a paired samples Wilcoxon test. Separate plots shown for: **(A)** Association with just a chloroplast. **(B)** Association with a chloroplast and the nucleus. **(C)** Association with just the nucleus.

**Figure S5:**
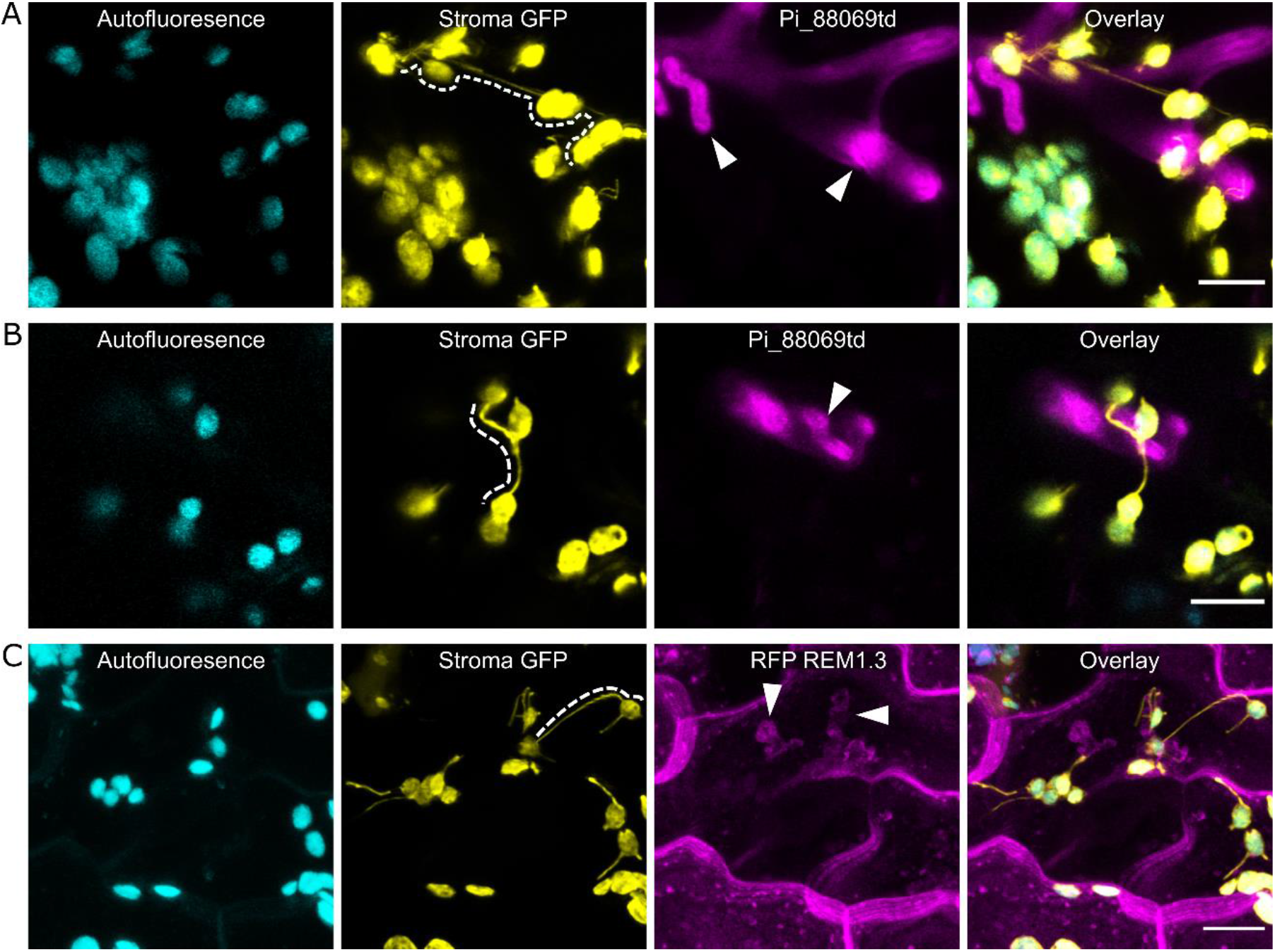
Stromules often bridge multiple chloroplasts, including those associated with different haustoria. Maximum projection confocal micrographs of *N. benthamiana* plants expressing GFP in chloroplast stroma (CpGFP) infected with *P. infestans* showing examples of stromules bridging multiple chloroplasts at haustoria. White dotted line shows the long distance stromule connection, white arrow heads mark haustoria. All scale bars are 10 μm. **(A-B)** Infection with *P. infestans* strain 88069td. **(C)** Infection with wild-type *P. infestans* and transient plant cell expression of RFP REM1.3 to mark the plasma membrane and extra haustorial membrane.

**Figure S6:**
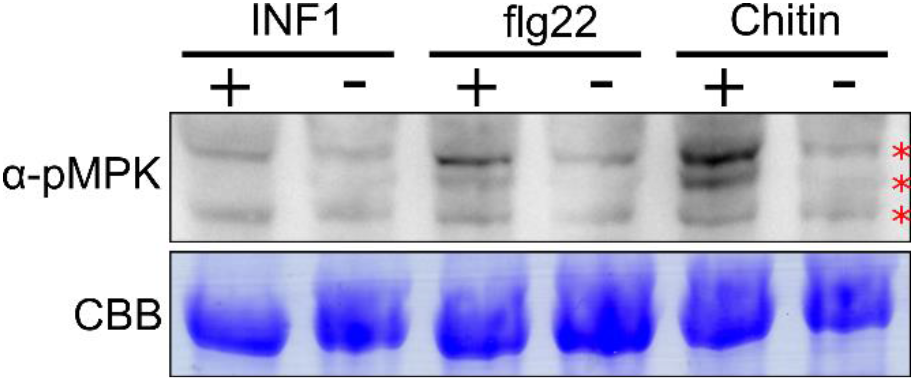
PAMPs induce MAPK phosphorylation at the concentrations shown to induce stromules. Western blot of tissue treated with INF1, flg22, chitin (+) or water (-) for 24 hr. Endogenous phosphorylated mitogen activated kinases detected at the expected sizes marker by red asterisks. Loading shown by Coomassie brilliant blue (CBB) staining.

**Figure S7:**
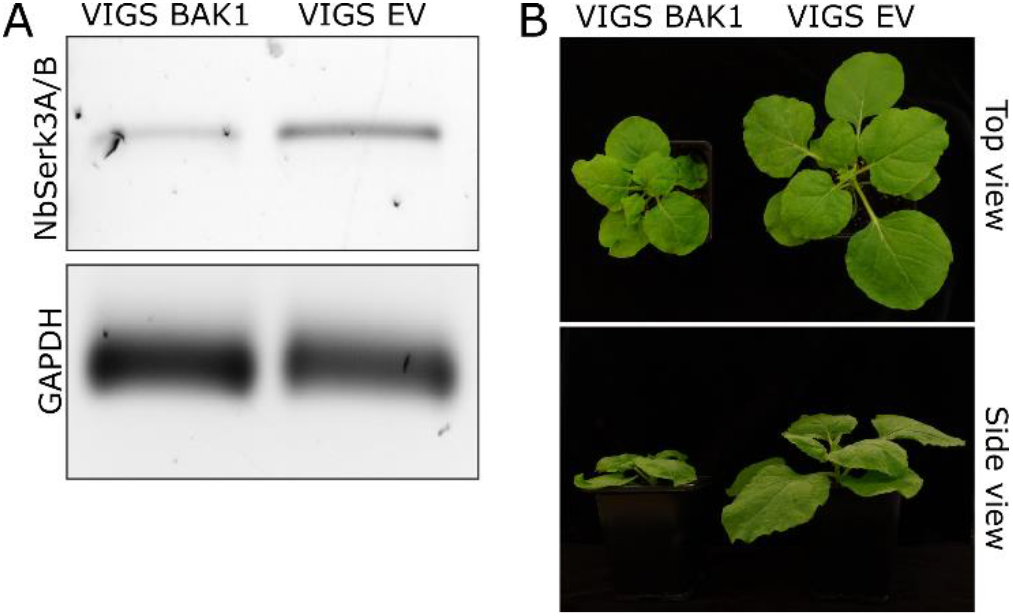
VIGS *BAK1* construct successfully reduces expression of *BAK1*. **(A)** Semi-quantitative RT-PCR of *BAK1* shows that it was silenced in VIGS *BAK1* tissue compared to EV control plants. RT-PCR of housekeeping GAPDH was used as an internal control for cDNA loading. For both, primers that did not amplify the silencing target were used. **(B)** Photos of representative *N. benthamiana* plants 5 weeks old, showing symptoms (notably stunting) of *BAK1* silencing by VIGS.

**Figure S8:**
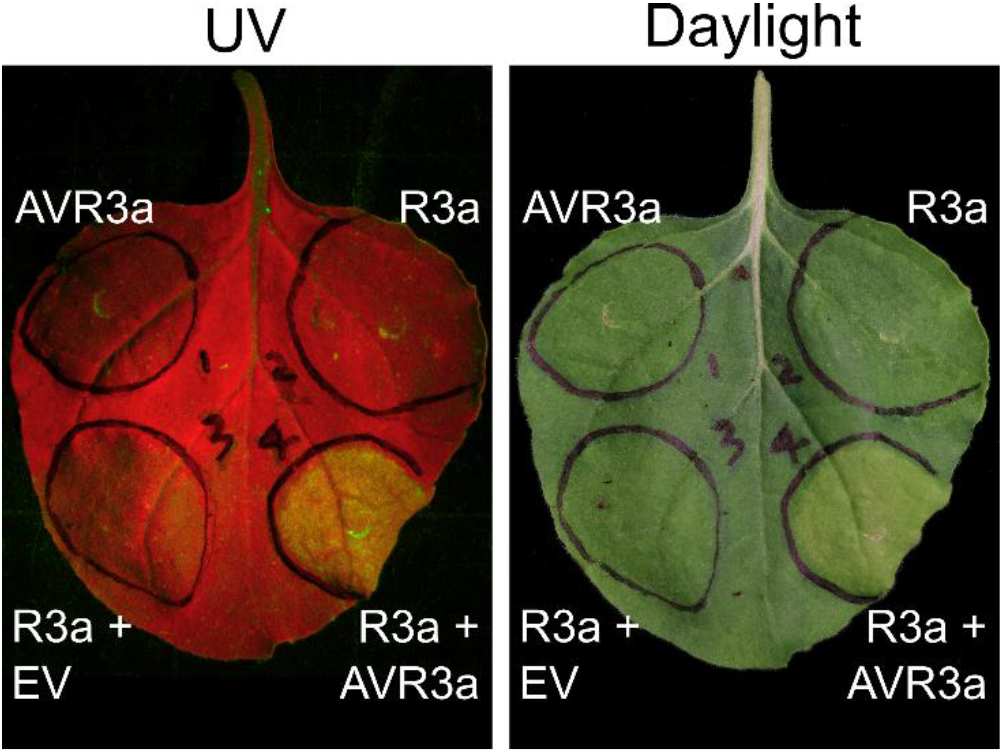
AVR3a induces cell-death in the presence of R3a. AVR3a or R3a were transiently expressed alone, triggering no cell death. EV and R3a also triggered no cell death. Only transient co-expression of AVR3a with R3a triggered cell death indicating functionality of the construct, as visible by white light or autofluorescence under UV light.

**Video S1: 3D image of Fig. 1A showing chloroplast focal accumulation at haustoria and stromules interacting with each other and other chloroplasts.** 3D visualization comprises Z-stack of confocal images of leaf epidermal cells from transplastomic CpGFP (Yellow) *N. benthamiana* plants infected with red-fluorescent *P. infestans* strain 88069td (Magenta).

**Video S2: Time-lapse series showing chloroplasts moving around a haustorium.** 3D time lapse visualization comprises Z-stack of confocal images of leaf epidermal cells from transplastomic CpGFP (Yellow) *N. benthamiana* plants infected with red-fluorescent *P. infestans* strain 88069td (Magenta).

**Video S3: 3D time-lapse series showing dynamic stromule interactions and relocation of chloroplasts towards a haustorium.** 3D visualization comprises Z-stack of confocal images of leaf epidermal cells from transplastomic CpGFP (Yellow) *N. benthamiana* plants infected with red-fluorescent *P. infestans* strain 88069td (Magenta).

**Video S4: Time-lapse series two chloroplasts navigating towards a haustorium without producing stromules.** Leaf epidermal cells from transplastomic CpGFP (Green) *N. benthamiana* plants red-fluorescent *P. infestans* strain 88069td (red), chlorophyll shown in blue. White arrowhead shows haustoria, purple arrow heads highlight the moving chloroplasts. Scale bar = 10 μm.

**Video S5: Time-lapse series showing chloroplasts accumulation to a haustorium.** Leaf epidermal cells from wild-type *N. benthamiana* plants expressing PM and EHM marker RFP:REM1.3 (Magenta) and endosomal marker GFP:RAB8 (Yellow) infected with wild-type *P. infestans* 88069. Blue is chlorophyll autofluorescence, labelling the chloroplasts.

**Video S6: Time-lapse series showing collapse of haustorium associated with a chloroplast.** Leaf epidermal cells from WT *N. benthamiana* plants expressing chloroplast outer envelope marker CHUP1:GFP (Yellow) and PM and EHM marker RFP:REM1.3 (Magenta) infected with wild-type *P. infestans* 88069. Blue is chlorophyll autofluorescence, labelling the chloroplasts. “H” indicates haustorium that collapses.

**Video S7: Time-lapse series showing collapse of haustorium associated with a chloroplast.** Leaf epidermal cells from WT *N. benthamiana* plants expressing chloroplast outer envelope marker CHUP1:GFP (Yellow) and PM and EHM marker RFP:REM1.3 (Magenta) infected with wild-type *P. infestans* 88069. Blue is chlorophyll autofluorescence, labelling the chloroplasts. “H” indicates haustorium that collapses.

**Video S8: Time-lapse series showing optical trapping of chloroplast in Fig. 1D which escapes the trap and springs back to the haustorium.** TIRF microscopy combined with laser capture in leaf epidermal cells from transplastomic CpGFP (channel not shown) *N. benthamiana* plants expressing PM and EHM marker RFP:REM1.3 (Grayscale) infected with wild-type *P. infestans* 88069. Chloroplast is visible due to chlorophyll autofluorescence overlapping with RFP emission spectrum. Scale bar is 10 μm.

**Video S9: Time-lapse series showing chloroplasts and stromules moving around a haustorium.** 3D time lapse visualization comprises Z-stack of confocal images of leaf epidermal cells from transplastomic CpGFP (Yellow) *N. benthamiana* plants infected with red-fluorescent *P. infestans* strain 88069td (Magenta). Grayscale crop of CpGFP signal highlights the extent of chloroplast stromules embracing the haustorium.

**Video S10: 3D image of Fig. S5A showing chloroplasts form long-distance stromule interactions that can bridge more than one haustorium.** 3D visualization comprises Z-stack of confocal images of leaf epidermal cells from transplastomic CpGFP (Yellow) *N. benthamiana* plants infected with red-fluorescent *P. infestans* strain 88069td (Magenta). ‘H’ indicates haustorium.

**Video S11: 3D image of chloroplast and stromules embracing a haustorium.** 3D visualization comprises Z-stack of confocal images of leaf epidermal cells from transplastomic CpGFP (Yellow) *N. benthamiana* plants expressing PM and EHM marker RFP:REM1.3 (Magenta) infected with wild-type *P. infestans* 88069.

**Video S12: Time-lapse series showing optical trapping of chloroplast in Fig. 3D and co-migration of a second chloroplast interacting via a stromule-like extension.** TIRF microscopy combined with laser capture in leaf epidermal cells from transplastomic CpGFP (Grayscale) *N. benthamiana* plants. Scale bar is 10 μm.

## Notes

### Competing Interest Statement

The authors have declared no competing interest.

### Summary of Updates

In V1 of the manuscript, we found an effect of silencing CHUP1 on chloroplast movement to haustoria. We now generated chup1 knockout lines and did not observe this phenotype. In V2 we have removed data relating to CHUP1 and will prepare a separate manuscript using chup1 knockout lines.

## References

Belhaj, K., Lin, B., & Mauch, F. (2009). The chloroplast protein RPH1 plays a role in the immune response of Arabidopsis to Phytophthora brassicae. Plant Journal, 58(2), 287–298. https://doi.org/10.1111/j.1365-313X.2008.03779.x

Belousov, V. V., Fradkov, A. F., Lukyanov, K. A., Staroverov, D. B., Shakhbazov, K. S., Terskikh, A. V., & Lukyanov, S. (2006). Genetically encoded fluorescent indicator for intracellular hydrogen peroxide. Nature Methods, 3(4), 281–286. https://doi.org/10.1038/nmeth866

Bozkurt, T. O., Belhaj, K., Dagdas, Y. F., Chaparro-Garcia, A., Wu, C. H., Cano, L. M., & Kamoun, S. (2015). Rerouting of Plant Late Endocytic Trafficking Toward a Pathogen Interface. Traffic, 16(2), 204–226. https://doi.org/10.1111/tra.12245

Bozkurt, T. O., Richardson, A., Dagdas, Y. F., Mongrand, S., Kamoun, S., & Raffaele, S. (2014). The plant membrane-associated REMORIN1.3 accumulates in discrete perihaustorial domains and enhances susceptibility to phytophthora infestans. Plant Physiology, 165(3), 1005–1018. https://doi.org/10.1104/pp.114.235804

Bozkurt, T. O., Schornack, S., Win, J., Shindo, T., Ilyas, M., Oliva, R., Cano, L. M., Jones, A. M. E., Huitema, E., Van Der Hoorn, R. A. L., & Kamoun, S. (2011). Phytophthora infestans effector AVRblb2 prevents secretion of a plant immune protease at the haustorial interface. Proceedings of the National Academy of Sciences of the United States of America, 108(51), 20832–20837. https://doi.org/10.1073/pnas.1112708109

Brunkard, J. O., Runkel, A. M., & Zambryski, P. C. (2015). Chloroplasts extend stromules independently and in response to internal redox signals. Proceedings of the National Academy of Sciences of the United States of America, 112(32), 10044–10049. https://doi.org/10.1073/pnas.1511570112

Caplan, J. L., Kumar, A. S., Park, E., Padmanabhan, M. S., Hoban, K., Modla, S., Czymmek, K., & Dinesh-Kumar, S. P. (2015). Chloroplast Stromules Function during Innate Immunity. Developmental Cell, 34(1), 45–57. https://doi.org/10.1016/j.devcel.2015.05.011

Chaparro-Garcia, A., Schwizer, S., Sklenar, J., Yoshida, K., Petre, B., Bos, J. I. B., Schornack, S., Jones, A. M. E., Bozkurt, T. O., & Kamoun, S. (2015). Phytophthora infestans RXLR-WY effector AVR3a associates with dynamin-related protein 2 required for endocytosis of the plant pattern recognition receptor FLS2. PLoS ONE, 10(9), e0137071. https://doi.org/10.1371/journal.pone.0137071

Chaparro-Garcia, A., Wilkinson, R. C., Gimenez-Ibanez, S., Findlay, K., Coffey, M. D., Zipfel, C., Rathjen, J. P., Kamoun, S., & Schornack, S. (2011). The receptor-like kinase serk3/bak1 is required for basal resistance against the late blight pathogen Phytophthora infestans in Nicotiana benthamiana. PLoS ONE, 6(1). https://doi.org/10.1371/journal.pone.0016608

Dagdas, Y. F., Pandey, P., Tumtas, Y., Sanguankiattichai, N., Belhaj, K., Duggan, C., Leary, A. Y., Segretin, M. E., Contreras, M. P., Savage, Z., Khandare, V. S., Kamoun, S., & Bozkurt, T. O. (2018). Host autophagy machinery is diverted to the pathogen interface to mediate focal defense responses against the irish potato famine pathogen. ELife, 7, 1–15. https://doi.org/10.7554/eLife.37476

Daniel, R., & Guest, D. (2006). Defence responses induced by potassium phosphonate in Phytophthora palmivora-challenged Arabidopsis thaliana. Physiological and Molecular Plant Pathology, 67(3-5), 194–201. https://doi.org/10.1016/j.pmpp.2006.01.003

Ding, X., Jimenez-Gongora, T., Krenz, B., & Lozano-Duran, R. (2019). Chloroplast clustering around the nucleus is a general response to pathogen perception in Nicotiana benthamiana. Molecular Plant Pathology, 20(9), 1298–1306. https://doi.org/10.1111/mpp.12840

Erickson, J. L., Adlung, N., Lampe, C., Bonas, U., & Schattat, M. H. (2018). The Xanthomonas effector XopL uncovers the role of microtubules in stromule extension and dynamics in Nicotiana benthamiana. Plant Journal, 93(5), 856–870. https://doi.org/10.1111/tpj.13813

Genre, A., Chabaud, M., Timmers, T., Bonfante, P., & Barker, D. G. (2005). Arbuscular mycorrhizal fungi elicit a novel intracellular apparatus in Medicago truncatula root epidermal cells before infection. Plant Cell, 17(12), 3489–3499. https://doi.org/10.1105/tpc.105.035410

Gray, J. C., Hansen, M. R., Shaw, D. J., Graham, K., Dale, R., Smallman, P., Natesan, S. K. A., & Newell, C. A. (2012). Plastid stromules are induced by stress treatments acting through abscisic acid. Plant Journal, 69(3), 387–398. https://doi.org/10.1111/j.1365-313X.2011.04800.x

Griffis, A. H. N., Groves, N. R., Zhou, X., & Meier, I. (2014). Nuclei in motion: Movement and positioning of plant nuclei in development, signaling, symbiosis, and disease. Frontiers in Plant Science, 5(APR), 1–7. https://doi.org/10.3389/fpls.2014.00129

Hanson, M. R., & Hines, K. M. (2018). Stromules: Probing formation and function. Plant Physiology, 176(1), 128–137. https://doi.org/10.1104/pp.17.01287

Heese, A., Hann, D. R., Gimenez-Ibanez, S., Jones, A. M. E., He, K., Li, J., Schroeder, J. I., Peck, S. C., & Rathjen, J. P. (2007). The receptor-like kinase SERK3/BAK1 is a central regulator of innate immunity in plants. Proceedings of the National Academy of Sciences of the United States of America, 104(29), 12217–12222. https://doi.org/10.1073/pnas.0705306104

Helle, S. C. J., Kanfer, G., Kolar, K., Lang, A., Michel, A. H., & Kornmann, B. (2013). Organization and function of membrane contact sites. In Biochimica et Biophysica Acta - Molecular Cell Research (Vol. 1833, Issue 11, pp. 2526–2541). Elsevier. https://doi.org/10.1016/j.bbamcr.2013.01.028

Hellens, R., Mullineaux, P., & Klee, H. (2000). A guide to Agrobacterium binary Ti vectors. Trends in Plant Science, 5(10), 446–451. https://doi.org/10.1016/S1360-1385(00)01740-4

Higa, T., Suetsugu, N., Kong, S. G., & Wada, M. (2014). Actin-dependent plastid movement is required for motive force generation in directional nuclear movement in plants. Proceedings of the National Academy of Sciences of the United States of America, 111(11), 4327–4331. https://doi.org/10.1073/pnas.1317902111

Hoitzing, H., Johnston, I. G., & Jones, N. S. (2015). What is the function of mitochondrial networks? A theoretical assessment of hypotheses and proposal for future research. BioEssays, 37(6), 687–700. https://doi.org/10.1002/bies.201400188

Jelenska, J., Yao, N., Vinatzer, B. A., Wright, C. M., Brodsky, J. L., & Greenberg, J. T. (2007). A J Domain Virulence Effector of Pseudomonas syringae Remodels Host Chloroplasts and Suppresses Defenses. Current Biology, 17(6), 499–508. https://doi.org/10.1016/j.cub.2007.02.028

Jones, J. D. G., & Dangl, J. L. (2006). The plant immune system. In Nature (Vol. 444, Issue 7117, pp. 323–329). Nature Publishing Group. https://doi.org/10.1038/nature05286

Kadota, A., Yamada, N., Suetsugu, N., Hirose, M., Saito, C., Shoda, K., Ichikawa, S., Kagawa, T., Nakano, A., & Wada, M. (2009). Short actin-based mechanism for light-directed chloroplast movement in Arabidopsis. Proceedings of the National Academy of Sciences of the United States of America, 106(31), 13106–13111. https://doi.org/10.1073/pnas.0906250106

Kamoun, S., Van West, P., Vleeshouwers, V. G. A. A., De Groot, K. E., & Govers, F. (1998). Resistance of Nicotiana benthamiana to Phytophthora infestans is mediated by the recognition of the elicitor protein INF1. Plant Cell, 10(9), 1413–1425. https://doi.org/10.1105/tpc.10.9.1413

Ketelaar, T., Meijer, H. J. G., Spiekerman, M., Weide, R., & Govers, F. (2012). Effects of latrunculin B on the actin cytoskeleton and hyphal growth in Phytophthora infestans. Fungal Genetics and Biology, 49(12), 1014–1022. https://doi.org/10.1016/j.fgb.2012.09.008

Kobayashi, I., & Hakuno, H. (2003). Actin-related defense mechanism to reject penetration attempt by a non-pathogen is maintained in tobacco BY-2 cells. Planta, 217(2), 340–345. https://doi.org/10.1007/s00425-003-1042-3

Koh, S., André, A., Edwards, H., Ehrhardt, D., & Somerville, S. (2005). Arabidopsis thaliana subcellular responses to compatible Erysiphe cichoracearum infections. Plant Journal, 44(3), 516–529. https://doi.org/10.1111/j.1365-313X.2005.02545.x

Kumar, A. S., Park, E., Nedo, A., Alqarni, A., Ren, L., Hoban, K., Modla, S., McDonald, J. H., Kambhamettu, C., Dinesh-Kumar, S. P., & Caplan, J. L. (2018). Stromule extension along microtubules coordinated with actin-mediated anchoring guides perinuclear chloroplast movement during innate immunity. ELife, 7, 1–33. https://doi.org/10.7554/eLife.23625

Kwon, C., Neu, C., Pajonk, S., Yun, H. S., Lipka, U., Humphry, M., Bau, S., Straus, M., Kwaaitaal, M., Rampelt, H., Kasmi, F. El, Jürgens, G., Parker, J., Panstruga, R., Lipka, V., & Schulze-Lefert, P. (2008). Co-option of a default secretory pathway for plant immune responses. Nature, 451(7180), 835–840. https://doi.org/10.1038/nature06545

Liu, L., & Li, J. (2019). Communications between the endoplasmic reticulum and other organelles during abiotic stress response in plants. In Frontiers in Plant Science (Vol. 10, p. 749). Frontiers Media S.A. https://doi.org/10.3389/fpls.2019.00749

Lukan, T., Županič, A., Mahkovec Povalej, T., Brunkard, J. O., Juteršek, M., Baebler, Š., & Gruden, K. (2021). Chloroplast redox state changes indicate cell-to-cell signalling during the hypersensitive response. BioRxiv.

Opalski, K. S., Schultheiss, H., Kogel, K. H., & Hückelhoven, R. (2005). The receptor-like MLO protein and the RAC/ROP family G-protein RACB modulate actin reorganization in barley attacked by the biotrophic powdery mildew fungus Blumeria graminis f.sp. hordei. Plant Journal, 41(2), 291–303. https://doi.org/10.1111/j.1365-313X.2004.02292.x

Padmanabhan, M. S., & Dinesh-Kumar, S. P. (2010). All hands on deck-the role of chloroplasts, endoplasmic reticulum, and the nucleus in driving plant innate immunity. Molecular Plant-Microbe Interactions, 23(11), 1368–1380. https://doi.org/10.1094/MPMI-05-10-0113

Pecrix, Y., Buendia, L., Penouilh-Suzette, C., Maréchaux, M., Legrand, L., Bouchez, O., Rengel, D., Gouzy, J., Cottret, L., Vear, F., & Godiard, L. (2019). Sunflower resistance to multiple downy mildew pathotypes revealed by recognition of conserved effectors of the oomycete Plasmopara halstedii. Plant Journal, 97(4), 730–748. https://doi.org/10.1111/tpj.14157

Petre, B., Lorrain, C., Saunders, D. G. O., Win, J., Sklenar, J., Duplessis, S., & Kamoun, S. (2016). Rust fungal effectors mimic host transit peptides to translocate into chloroplasts. Cellular Microbiology, 18(4), 453–465. https://doi.org/10.1111/cmi.12530

Rocchetti, A., Hawes, C., & Kriechbaumer, V. (2014). Fluorescent labelling of the actin cytoskeleton in plants using a cameloid antibody. Plant Methods, 10(1), 12. https://doi.org/10.1186/1746-4811-10-12

Schattat, M., Barton, K., Baudisch, B., Klösgen, R. B., & Mathur, J. (2011). Plastid stromule branching coincides with contiguous endoplasmic reticulum dynamics. Plant Physiology, 155(4), 1667–1677. https://doi.org/10.1104/pp.110.170480

Schattat, M. H., Barton, K. A., & Mathur, J. (2015). The myth of interconnected plastids and related phenomena. Protoplasma, 252(1), 359–371. https://doi.org/10.1007/s00709-014-0666-4

Scheler, B., Schnepf, V., Galgenmüller, C., Ranf, S., & Hückelhoven, R. (2016). Barley disease susceptibility factor RACB acts in epidermal cell polarity and positioning of the nucleus. Journal of Experimental Botany, 67(11), 3263–3275. https://doi.org/10.1093/jxb/erw141

Schmelzer, E. (2002). Cell polarization, a crucial process in fungal defence. Trends in Plant Science, 7(9), 411–415. https://doi.org/10.1016/S1360-1385(02)02307-5

Schneider, C. A., Rasband, W. S., & Eliceiri, K. W. (2012). NIH Image to ImageJ: 25 years of image analysis. In Nature Methods (Vol. 9, Issue 7, pp. 671–675). Nature Publishing Group. https://doi.org/10.1038/nmeth.2089

Silva, B. S. C., DiGiovanni, L., Kumar, R., Carmichael, R. E., Kim, P. K., & Schrader, M. (2020). Maintaining social contacts: The physiological relevance of organelle interactions. In Biochimica et Biophysica Acta - Molecular Cell Research (Vol. 1867, Issue 11, p. 118800). Elsevier B.V. https://doi.org/10.1016/j.bbamcr.2020.118800

Song, J., Win, J., Tian, M., Schornack, S., Kaschani, F., Ilyas, M., Van Der Hoorn, R. A. L., & Kamoun, S. (2009). Apoplastic effectors secreted by two unrelated eukaryotic plant pathogens target the tomato defense protease Rcr3. Proceedings of the National Academy of Sciences of the United States of America, 106(5), 1654–1659. https://doi.org/10.1073/pnas.0809201106

Sparkes, I., White, R. R., Coles, B., Botchway, S. W., & Ward, A. (2018). Using optical tweezers combined with total internal reflection microscopy to study interactions between the ER and Golgi in plant cells. In Methods in Molecular Biology (Vol. 1691, pp. 167–178). Humana Press Inc. https://doi.org/10.1007/978-1-4939-7389-7_13

Stegemann, S., Keuthe, M., Greiner, S., & Bock, R. (2012). Horizontal transfer of chloroplast genomes between plant species. Proceedings of the National Academy of Sciences of the United States of America, 109(7), 2434–2438. https://doi.org/10.1073/pnas.1114076109

Su, J., Yang, L., Zhu, Q., Wu, H., He, Y., Liu, Y., Xu, J., Jiang, D., & Zhang, S. (2018). Active photosynthetic inhibition mediated by MPK3/MPK6 is critical to effector-triggered immunity. PLoS Biology, 16(5), 1–29. https://doi.org/10.1371/journal.pbio.2004122

Suetsugu, N., Higa, T., Gotoh, E., & Wada, M. (2016). Light-induced movements of chloroplasts and nuclei are regulated in both cp-actin-filament-dependent and - independent manners in arabidopsis thaliana. PLoS ONE, 11(6), e0157429. https://doi.org/10.1371/journal.pone.0157429

Suetsugu, N., & Wada, M. (2016). Evolution of the Cp-Actin-based motility system of chloroplasts in green plants. Frontiers in Plant Science, 7(MAY2016), 561. https://doi.org/10.3389/fpls.2016.00561

Tang, C., Deng, L., Chang, D., Chen, S., Wang, X., & Kang, Z. (2016). TaADF3, an Actin-Depolymerizing factor, negatively modulates wheat resistance against puccinia striiformis. Frontiers in Plant Science, 6(JAN2016), 1–14. https://doi.org/10.3389/fpls.2015.01214

Van Damme, M., Zeilmaker, T., Elberse, J., Andel, A., De Sain-van Der Velden, M., & Van Den Ackerveken, G. (2009). Downy mildew resistance in arabidopsis by mutation of Homoserine Kinase. Plant Cell, 21(7), 2179–2189. https://doi.org/10.1105/tpc.109.066811

Van West, P., De Jong, A. J., Judelson, H. S., Emons, A. M. C., & Govers, F. (1998). The ipiO gene of phytophthora infestans is highly expressed in invading hyphae during infection. Fungal Genetics and Biology, 23(2), 126–138. https://doi.org/10.1006/fgbi.1998.1036

Wada, M., & Kong, S. G. (2018). Actin-mediated movement of chloroplasts. In Journal of Cell Science (Vol. 131, Issue 2). Company of Biologists Ltd. https://doi.org/10.1242/jcs.210310

Wang, S., Boevink, P. C., Welsh, L., Zhang, R., Whisson, S. C., & Birch, P. R. J. (2017). Delivery of cytoplasmic and apoplastic effectors from Phytophthora infestans haustoria by distinct secretion pathways. New Phytologist. https://doi.org/10.1111/nph.14696

Whisson, S. C., Boevink, P. C., Moleleki, L., Avrova, A. O., Morales, J. G., Gilroy, E. M., Armstrong, M. R., Grouffaud, S., Van West, P., Chapman, S., Hein, I., Toth, I. K., Pritchard, L., & Birch, P. R. J. (2007). A translocation signal for delivery of oomycete effector proteins into host plant cells. Nature, 450(7166), 115–118. https://doi.org/10.1038/nature06203

Whisson, S. C., Boevink, P. C., Wang, S., & Birch, P. R. (2016). The cell biology of late blight disease. Current Opinion in Microbiology, 34, 127–135. https://doi.org/10.1016/j.mib.2016.09.002

Wu, C. H., Abd-El-Haliem, A., Bozkurt, T. O., Belhaj, K., Terauchi, R., Vossen, J. H., & Kamoun, S. (2017). NLR network mediates immunity to diverse plant pathogens. Proceedings of the National Academy of Sciences of the United States of America, 114(30), 8113–8118. https://doi.org/10.1073/pnas.1702041114

Zabala, M. de T., Littlejohn, G., Jayaraman, S., Studholme, D., Bailey, T., Lawson, T., Tillich, M., Licht, D., Bölter, B., Delfino, L., Truman, W., Mansfield, J., Smirnoff, N., & Grant, M. (2015). Chloroplasts play a central role in plant defence and are targeted by pathogen effectors. Nature Plants, 1(6), 15074. https://doi.org/10.1038/NPLANTS.2015.74

